# Nanopore sequencing and assembly of a human genome with ultra-long reads

**DOI:** 10.1101/128835

**Authors:** Miten Jain, S Koren, J Quick, AC Rand, TA Sasani, JR Tyson, AD Beggs, AT Dilthey, IT Fiddes, S Malla, H Marriott, KH Miga, T Nieto, J O’Grady, HE Olsen, BS Pedersen, A Rhie, H Richardson, AR Quinlan, TP Snutch, L Tee, B Paten, AM Phillippy, JT Simpson, NJ Loman, M Loose

**Affiliations:** UC Santa Cruz Genomics Institute, University of California, Santa Cruz, CA, USA; Genome Informatics Section, Computational and Statistical Genomics Branch, National Human Genome Research Institute, Bethesda, Maryland, USA; Institute of Microbiology and Infection, University of Birmingham, Birmingham B15 2TT, UK; Department of Human Genetics, University of Utah, Salt Lake City, UT, USA; USTAR Center for Genetic Discovery, University of Utah, Salt Lake City, UT, USA; Department of Biomedical Informatics, University of Utah, Salt Lake City, UT, USA; Michael Smith Laboratories and Djavad Mowafaghian Centre for Brain Health, University of British Columbia, Vancouver, Canada; Surgical Research Laboratory, Institute of Cancer & Genomic Science, University of Birmingham, UK; DeepSeq, School of Life Sciences, University of Nottingham, UK; Norwich Medical School, University of East Anglia, Norwich, UK; Ontario Institute for Cancer Research, Toronto M5G 0A3, Canada; Department of Computer Science, University of Toronto, Toronto M5S 3G4, Canada

## Abstract

Nanopore sequencing is a promising technique for genome sequencing due to its portability, ability to sequence long reads from single molecules, and to simultaneously assay DNA methylation. However until recently nanopore sequencing has been mainly applied to small genomes, due to the limited output attainable. We present nanopore sequencing and assembly of the GM12878 Utah/Ceph human reference genome generated using the Oxford Nanopore MinION and R9.4 version chemistry. We generated 91.2 Gb of sequence data (∼30× theoretical coverage) from 39 flowcells. *De novo* assembly yielded a highly complete and contiguous assembly (NG50 ∼3Mb). We observed considerable variability in homopolymeric tract resolution between different basecallers. The data permitted sensitive detection of both large structural variants and epigenetic modifications. Further we developed a new approach exploiting the long-read capability of this system and found that adding an additional 5×-coverage of ‘ultra-long’ reads (read N50 of 99.7kb) more than doubled the assembly contiguity. Modelling the repeat structure of the human genome predicts extraordinarily contiguous assemblies may be possible using nanopore reads alone. Portable *de novo* sequencing of human genomes may be important for rapid point-of-care diagnosis of rare genetic diseases and cancer, and monitoring of cancer progression. The complete dataset including raw signal is available as an Amazon Web Services Open Dataset at: https://github.com/nanopore-wgs-consortium/NA12878.

## Introduction

The human genome is a yardstick for assessing performance of DNA sequencing instruments ^1–5^. Despite continuous technical improvements, it remains challenging to sequence a human genome to high accuracy and completeness. This is due to its large size (a euchromatic genome of ∼3.1 Gb), heterozygosity, regions of extreme GC% bias, and because nearly half the genome is comprised of diverse families of repeats and large segmental duplications that range up to 1.7 Mbp in size ^6^. The repetitive structure poses major challenges for *de novo* assembly with extant “short read” sequencing technologies (∼25–300 bp for Illumina and ∼100–400 bp for Ion Torrent). Data generated by these instruments, whilst enabling highly accurate genotyping in non-repetitive regions, do not provide contiguous *de novo* assemblies nor a true map of the genome. This limits their ability to reconstruct repetitive sequences, detect complex structural variation, and fully characterize the genomes of other organisms that lack completed reference genomes.

Single-molecule sequencers, particularly those developed by Pacific Biosciences, can produce average read lengths of 10 kb or higher, making *de novo* assembly more tractable ^7^. In May 2014, a new single molecule sequencer, the MinION (Oxford Nanopore Technologies, UK) was made available to early access users ^8^. The MinION instrument is notable for its low capital cost ($1000 setup fee) and pocket-size portability. Until recently, the MinION has mainly been used for sequencing microbial genomes or PCR products ^9,10^, as the output of the instrument has been relatively low (up to 2 Gb, but typically <500 Mb). However, the potential of nanopore sequencing to enable contiguous de novo assemblies was previously demonstrated on the *Escherichia coli* K-12 genome, which was assembled into a single circular sequence using nanopore reads alone ^11^. More recently, assemblies of small eukaryotic genomes including yeasts, fungi and *C. elegans* have been demonstrated ^12–14^.

Recent updates to the integrated protein pore (a laboratory-evolved mutant of *E. coli* CsgG named R9.4), new library preparation techniques (1D ligation and 1D rapid), increases to sequencing speed (450 bases/s), and updated control software have resulted in much improved sequencing throughput making whole human genome sequencing feasible ^14–16^. Here we present the sequencing and assembly of a reference standard human genome, GM12878 from the Utah/CEPH pedigree using the MinION R9.4 1D chemistry, including ultra-long reads up to 882 kb in length. This genome was selected because it has been sequenced on a wide variety of platforms and has well validated variation call sets for performance benchmarking ^17^.

## Results

### Summary of dataset

Five individual laboratories collaborated to sequence human genomic DNA from the GM12878 human cell line. All DNA was sequenced directly, thereby avoiding PCR and preserving epigenetic modifications such as DNA methylation. In total, 39 MinION flowcells generated 14,183,584 reads containing 91,240,120,433 bases with a read N50 of 10,589 bp (SI Tables 1–4). This includes only reads generated by standard protocols and does not include ultra-long reads, which are described separately below. Sequencing read length was dependent on the input DNA (Figure 1A). Average yield per flow cell (2.3 Gb) was unrelated to DNA preparation methods (Figure 1B). 94.15% of reads had at least one alignment to the human reference (GRCh38) and 74.49% had a single alignment covering over 90% of their length. Median coverage depth was 26 fold and 96.95% (3.01/3.10 Gbp) bases of the reference were covered by at least one read (Figure 1C). The median identity of these reads was 84.06% (82.73% mean, 5.37% standard deviation). Similar to other single-molecule sequencing technologies, no length-bias was observed in the error rate with the MinION (Figure 1D).

**Figure 1.**
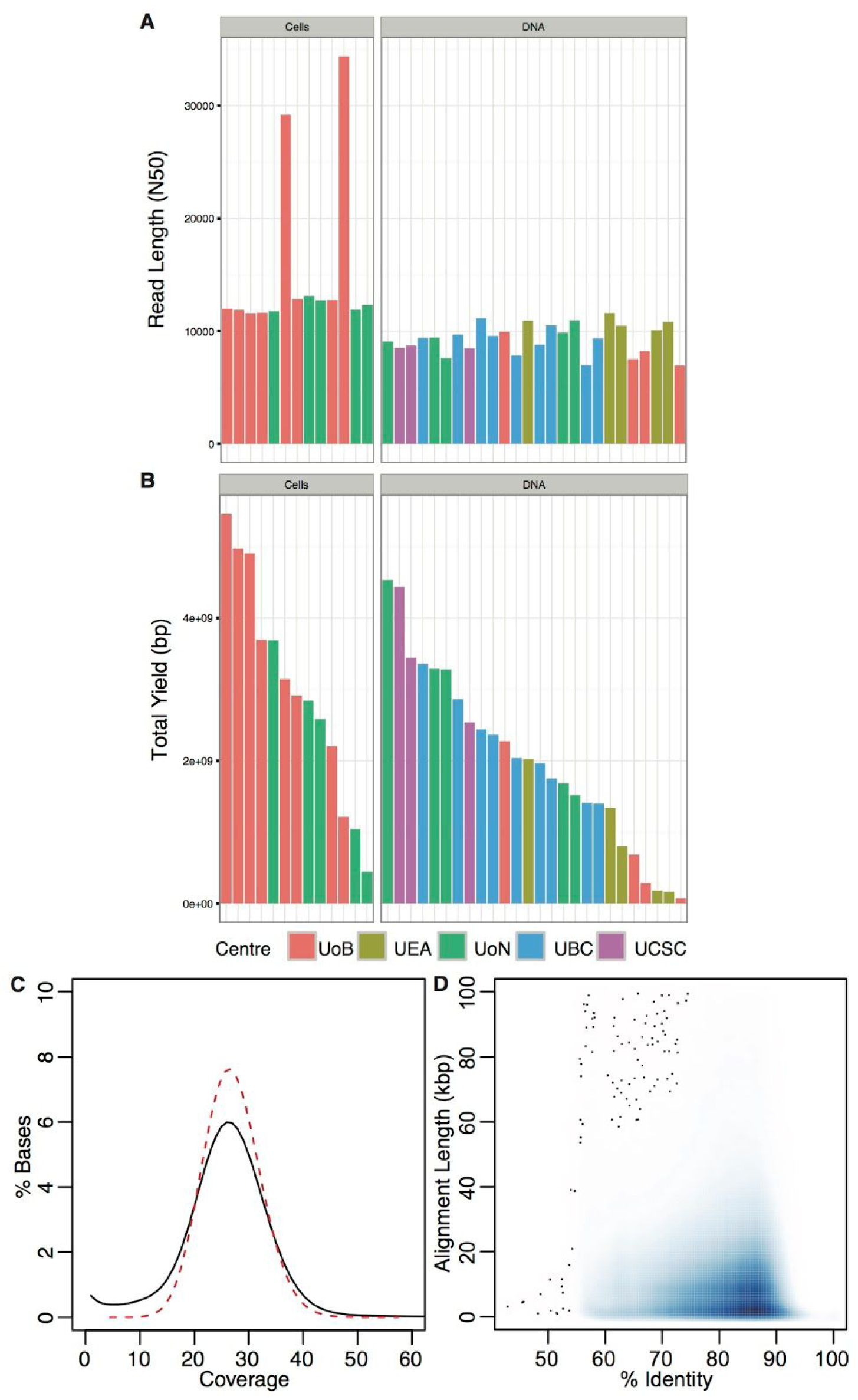
Summary of Dataset. A) Read Length N50s by flow cell coloured by sequencing center. Cells: DNA extracted directly from cell culture. DNA: Pre-extracted DNA purchased from Coriell. B) Total yield per flow cell grouped as A. C) Coverage of GRCh38 reference compared to a Poisson distribution. Reads were aligned to the 1000 genome GRCh38 reference. The depth of coverage of each reference position was tabulated using samtools depth and compared with a Poisson distribution with lambda = 27.4 (dashed red line) (mean coverage excluding 0-coverage positions). The true coverage distribution matches the expected Poisson distribution D) Alignment identity compared to alignment length. Alignments covering >90% of a sequence were extracted from bam files and identity calculated. The identity versus alignment length was plotted. The majority of alignments are between 80 and 90% identity and <20 Kb. No length bias was observed, with long alignments having the same identity as short ones.

### Sequence analysis and base caller evaluation

The base-calling algorithm used to decode raw ionic current signal can influence the resultant sequence calls. To characterize this effect, we selected reads mapping to chromosome 20 and performed base-calling using three methods available from Oxford Nanopore Technologies: the Metrichor cloud-based service; Nanonet, an open-source recurrent neural network (RNN); and Scrappie, a transducer neural network. Of note, we observed that a substantial fraction of the Scrappie output (4.7% reads, 14% bases) was composed of low-complexity sequence (SI Figure 1), which was removed before downstream analysis.

To assess nanopore sequencing read accuracy we realigned reads using a trained alignment model as previously described ^18^. Briefly, alignments generated by BWA-MEM were chained such that each read has at most one maximal alignment to the reference sequence (scored by length). The chained alignments were then used to derive the maximum likelihood estimate of alignment model parameters ^19^, and the trained model used to realign the reads. The median identity after re-alignment for Metrichor, Nanonet, and Scrappie base-called reads was 82.43%, 85.50%, and 86.05%, respectively. In chained alignments wherein the model was not used we observed a purine-to-purine substitution bias (SI Figure 2). The alignments produced by the trained model showed an improved substitution error rate, decreasing the overall transversion rate, albeit transition errors remained dominant.

To measure potential bias at the *k*-mer level, we compared counts of 5-mers in reads derived from chromosome 20. In Metrichor reads, the most underrepresented 5-mers were A/T-rich homopolymers. The most over-represented *k*-mers were G/C-rich and non-homopolymeric (SI Table 5). In contrast, Scrappie showed no underrepresentation of homopolymeric 5-mers and had a slight over representation of A/T homopolymers. Overall, Scrappie showed the lowest *k*-mer representation bias (Figure 2A). The improved homopolymer resolution of Scrappie was confirmed by inspection of chromosome 20 homopolymer calls versus the human reference (Figure 2B, SI Figure 3, Methods) ^20^. Despite this reduced bias, whole-genome assembly and analyses proceeded with the Metrichor reads, since Scrappie was in early development at the time of writing.

**Figure 2.**
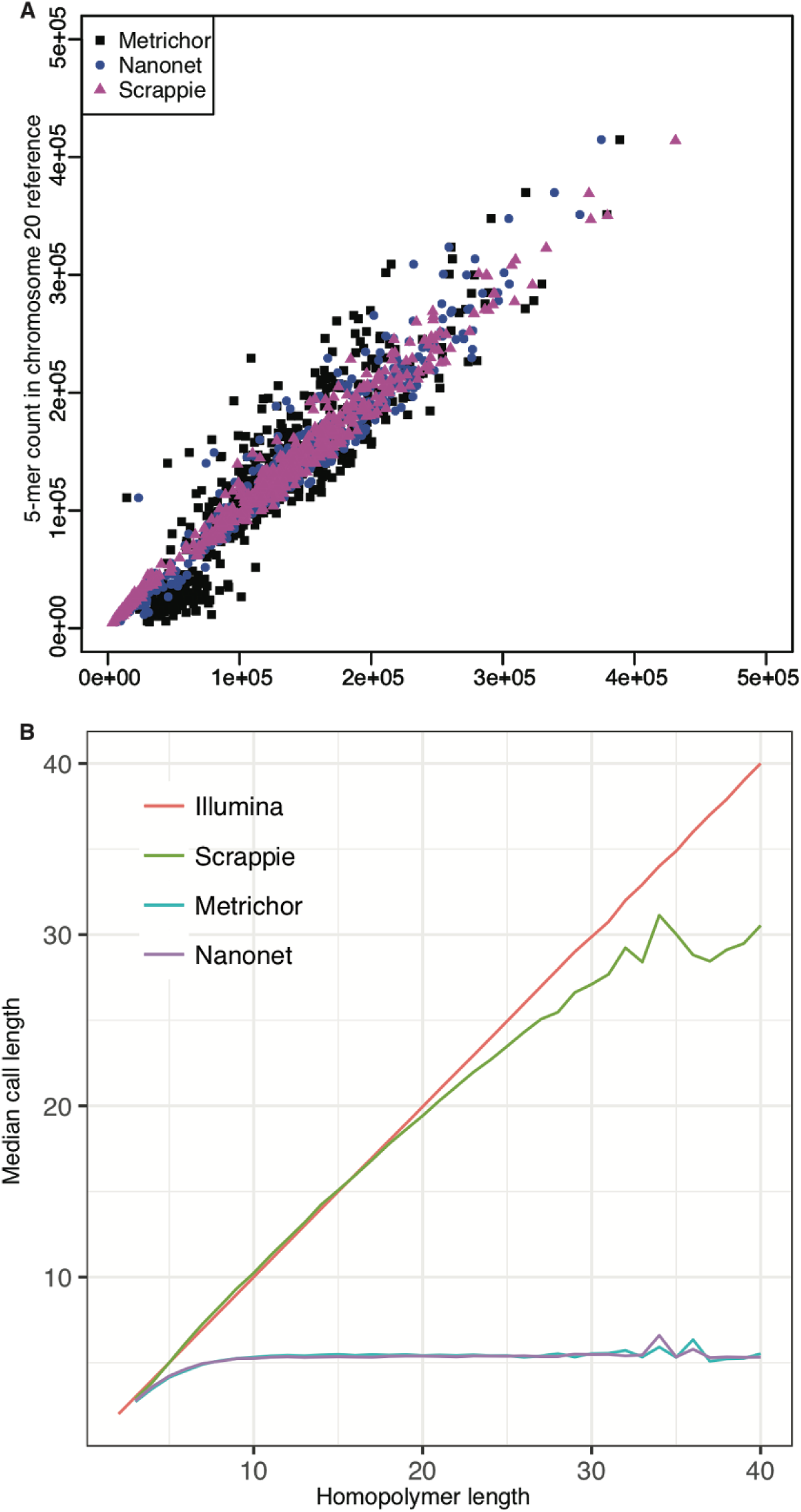
Base Caller Comparison and Homopolymer Resolution. A) Correlation between 5-mer counts in reads produced by various base callers compared to expected counts in the chromosome 20 reference. B) Chromosome 20 homopolymer length versus median homopolymer base-call length measured from individual Illumina and nanopore reads (Scrappie, Metrichor, and Nanonet). Metrichor and Nanonet base callers fail to produce homopolymer runs longer than ∼5 bp, resulting in a large deletion bias. Scrappie shows better correlation for longer homopolymer runs, but tends to over-call short homopolymers (between 5 and 15 bp) and under-call long homopolymers (>15 bp). Plot noise for longer homopolymers is due to fewer samples available at that length.

### Assembly of the dataset

We performed a *de novo* assembly of the 30× dataset with Canu ^21^ and polished the assembly using both nanopore signal and Illumina data (Table 1). Our initial assembly comprised 2,886 contigs with an NG50 contig size of 3 Mbp (NG50, the longest contig such that contigs of this length or greater sum to at least half the haploid genome size). We aligned the assembled contigs to the GRCh38 reference demonstrating agreement with previous GM12878 assemblies (SI Figure 4) ^22^. The number of identified structural differences (899) was similar to a previously published PacBio assembly of GM12878 (692) and comparable to other human genome assemblies ^5,21^, but with a higher than expected number of deletions due to consistent truncation of homopolymer and low-complexity regions (SI Figure 5, SI Table 6). Consensus identity versus GRCh38 was estimated to be 95.20% (Table 1). However, since GRCh38 is a composite of multiple human haplotypes, this represents a lower bound on accuracy. Comparisons against independent Illumina data from GM12878 yielded a slightly higher accuracy estimate of 95.74%.

**Table 1.** Summary Assembly Statistics.

| Assembly | Polishing | Contigs | # Bases (Mbp) | Max Contig (kb) | NG50 (kb) | GRCh38 Identity | GM12878 Identity |
| --- | --- | --- | --- | --- | --- | --- | --- |
| WGS Metrichor | N/A | 2,886 | 2,646.01 | 27,160 | 2,964 | 95.20% | 95.74% |
|  | Pilon x2 |  | 2,763.18 | 28,413 | 3,206 | 99.29% | 99.88% |
| Chr 20 Metrichor | N/A | 85 | 57.83 | 7,393 | 3,047 | 94.90% | 95.50% |
|  | Nanopolish |  | 60.35 | 7,667 | 5,394 | 98.84% | 99.24% |
|  | Pilon x2 |  | 60.58 | 7,680 | 5,423 | 99.33% | 99.89% |
|  | Nano + Pilon x2 |  | 60.76 | 7,698 | 5,435 | 99.64% | 99.95% |
| Chr 20 Nanonet | N/A | 74 | 58.51 | 12,314 | 2,849 | 95.99% | 96.64% |
|  | Nanopolish |  | 60.33 | 12,664 | 2,941 | 98.85% | 99.23% |
|  | Pilon x2 |  | 60.59 | 12,723 | 2,959 | 99.41% | 99.92% |
|  | Nano + Pilon x2 |  | 60.74 | 12,747 | 2,967 | 99.65% | 99.96% |
| Chr 20 Scrappie | N/A | 74 | 59.39 | 8,415 | 2,643 | 97.43% | 97.80% |
|  | Nanopolish |  | 60.15 | 8,521 | 2,681 | 99.12% | 99.44% |
|  | Pilon x2 |  | 60.36 | 8,541 | 2,691 | 99.64% | 99.95% |
|  | Nano + Pilon x2 |  | 60.34 | 8,545 | 2,691 | 99.70% | 99.96% |

Summary of assembly statistics. Whole genome assembly (WGS) was performed with reads base called by Metrichor. Chromosome 20 was assembled with reads produced by Metrichor and two additional base calling algorithms. All datasets contained 30× coverage of the genome/chromosome. The GRCh38 identities were computed based on 1-1 alignments to the GRCh38 reference including alt sites. A GM12878 reference was estimated using an independent sequencing dataset<sup>17</sup> (Methods).

Despite the low consensus accuracy, contiguity was good. For example, the assembly included a single ∼3 Mbp contig spanning all class I HLA genes from the major histocompatibility complex (MHC) region on chromosome 6, a region notoriously difficult to assemble from short reads. The more repetitive region of the MHC, containing class II HLA genes, was more fragmented but most genes were recovered in a single contig. Overall, the classical class I and II typing genes were successfully matched against a database of known alleles to infer HLA types, albeit with some error in the assembly (Methods, SI Table 7). In this highly diverse region of the genome, haplotype switching was evident in the Canu contigs, but phasing appears possible. As an example, a small part of the class II region is covered by two contigs, suggesting assembly of the second haplotype. For the larger collapsed contig, spanning most of the class II genes, heterozygous sites could be completely phased using the nanopore reads (SI Figure 6).

To improve base-accuracy of the initial assembly we mapped whole-genome Illumina (SRA:ERP001229) data to the contigs for polishing. Using Pilon we improved the estimated accuracy of the assembly to 99.88% (Table 1, SI Figure 7) ^23^. However limitations in mapping short Illumina reads to repetitive regions means the most repetitive areas of the assembly could not be corrected.

To further evaluate these assemblies, comparative annotation was performed both before and after polishing. This process used whole-genome alignments to project annotations from the GRCh38 reference and a combination of tools to clean the alignments and produce an annotation set (Methods). 58,338 genes (19,436 coding / 96.4% of genes in GENCODE V24 / 98.2% of coding genes) were identified representing 179,038 transcripts in the polished assembly. Reflecting the assembly’s high contiguity, only 857 (0.1%) of genes were found on two or more contigs.

Alternative approaches to improve assembly accuracy, using several different base-callers and exploiting signal-level nanopore data, were attempted on the subset of reads mapping to chromosome 20. To quantify the effect of base-calling on the assembly, each read set was re-assembled with the same Canu parameters used for the whole-genome dataset. Whilst all assemblies had similar contiguity, the assembly of the Scrappie reads improved accuracy from 95.74% to 97.80%. Further signal-level polishing using nanopolish increased accuracy to 99.44%, the highest accuracy achieved from nanopore data alone. Combining with Illumina data reached an accuracy of 99.96% (Table 1).

### Analysis of sequences not included in the primary assembly

To better understand sequences omitted from the primary genome analysis we assessed 1,425 degenerate contigs (26 Mbp) and corrected reads not incorporated into contigs (10.4 Gbp). The majority of sequences represented particular repeat classes e.g. LINEs, SINEs etc., as described in SI Figure 8. These were observed in similar proportion in the primary assembly, with the exception of satellite DNAs known to be enriched in human centromeric regions. Such satellites were enriched 2.93× in the unassembled data and 7.9× in the degenerate contigs. Additional sequence characterization at each centromeric transition in the primary Canu assembly determined that the majority of assembled centromeric satellites were present as individual contigs, with the largest assembled satellite assembly defined by a 94 kbp tandem repeat specific to centromere 15 (D15Z1, tig00007244).

### Measuring SNP and SV genotyping sensitivity

Using SVTyper, a Bayesian structural variant (SV) genotyper ^24^, we genotyped 2,435 previously identified GM12878 SVs ^25^ using Platinum Illumina WGS alignments. We re-examined these genotypes with nanopore alignments and a modified version of SVTyper (Methods). By measuring the concordance of Illumina and nanopore-derived genotypes at each site, we determined the sensitivity of SV genotyping as a function of the number of flowcells utilized (Figure 3A). Using all 39 flowcells, nanopore data recovered approximately 93% of high-confidence SVs with a false-positive rate of approximately 6% (Methods). Illumina and nanopore genotypes agreed at 82% of heterozygous and 91% of homozygous alternate sites.

**Figure 3.**
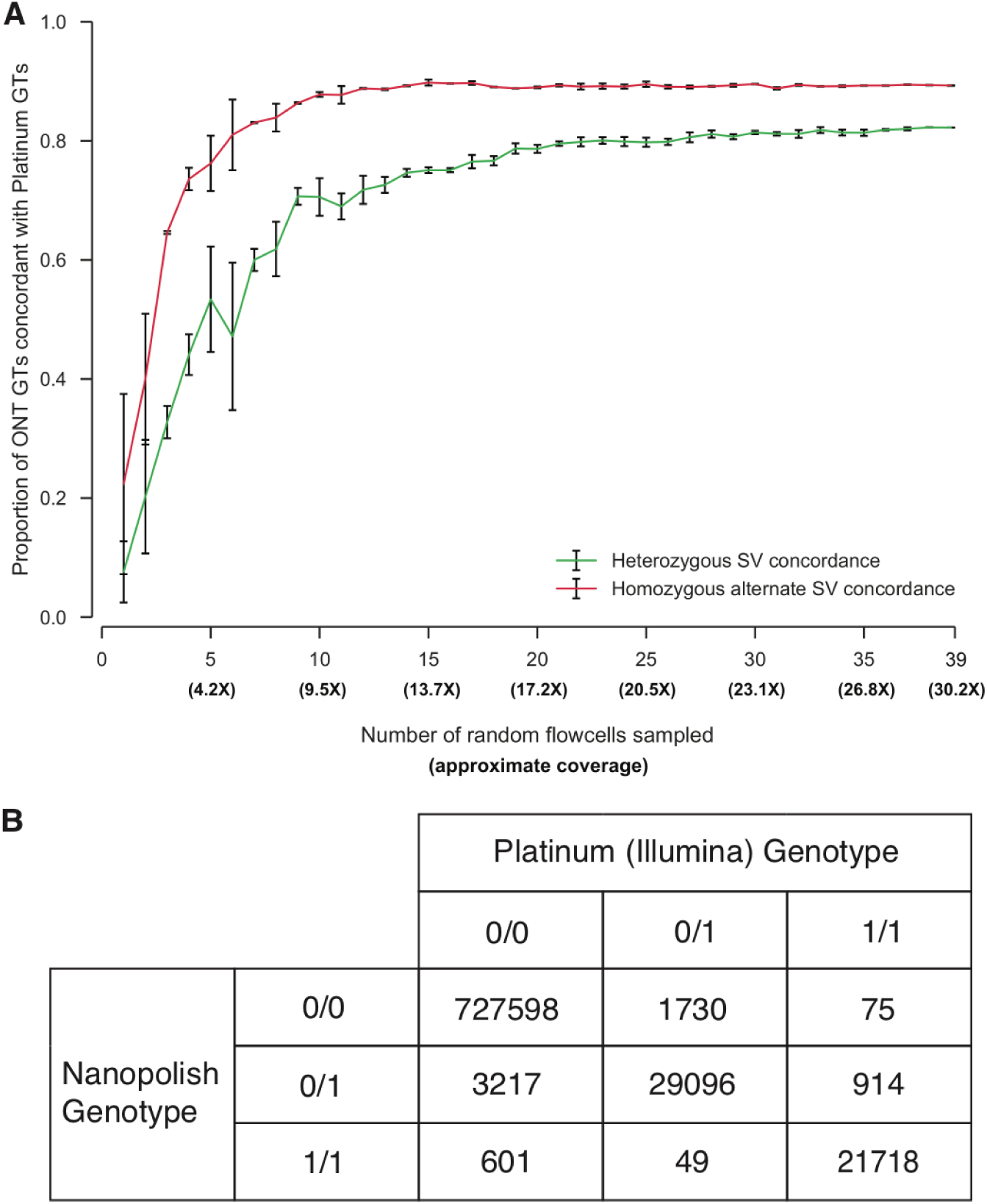
Structural Variation and Genotyping. A) Structural variant genotyping sensitivity using ONT reads. Genotypes were inferred for a set of 2,435 SVs using both Oxford Nanopore and Platinum Genomes (Illumina) alignments. Using reads from a random sample of *X* flowcells, sensitivity was calculated as the proportion of ONT-derived genotypes that were concordant with Illumina-derived genotypes. Error bars represent the standard deviation of sensitivity measurements from three randomly sampled sets of flowcells. B) Confusion matrix for genotype calling evaluation. Each cell contains the number of 1000 Genome sites for a particular (nanopolish, platinum) genotype combination.

We evaluated nanopore data for calling genotypes at known single nucleotide polymorphisms (SNPs) using signal-level data by calling genotypes at non-singleton SNPs on chromosome 20 from phase 3 of the 1000 Genomes ^26^ (Methods) and comparing these calls to Illumina platinum calls. The results are summarized as a confusion matrix (Figure 3B). A total of 99.16% of genotype calls are correct (778,412 out of 784,998 sites). This result is dominated by the large number of homozygous reference sites. If we assess accuracy by the fraction of called variant sites (heterozygous or homozygous non-reference) that have the correct genotype, the accuracy of our caller is 91.40% (50,814 out of 55,595), with the predominant error being mis-calling a site that should be homozygous reference as heterozygous (3,217 errors). Genotype accuracy when performing the reverse comparison, at sites annotated as variants in the platinum call set, is 94.83% (50,814 correct out of 53,582).

### Native 5-methyl cytosine detection

Nanopore sequencing detects DNA modifications as subtle changes to the ionic current when compared to unmodified bases ^27,28^. We employed two recently published algorithms, nanopolish and SignalAlign, to map 5-methyl cytosine at CpG dinucleotides on chromosome 20 of the GRCh38 reference ^29,30^. Briefly, the models align the ionic current readout to a reference sequence to infer the methylation status of a given base. Nanopolish outputs a frequency of reads calling a methylated cytosine and SignalAlign outputs a marginal probability of methylation summed over reads. We compared the output of both methods to published bisulfite data (ENCFF835NTC). Overall we observed good concordance with the published bisulfite sequencing results; the r-values for nanopolish and SignalAlign were 0.895 and 0.779 respectively (Figure 4, SI Figure 9).

**Figure 4.**
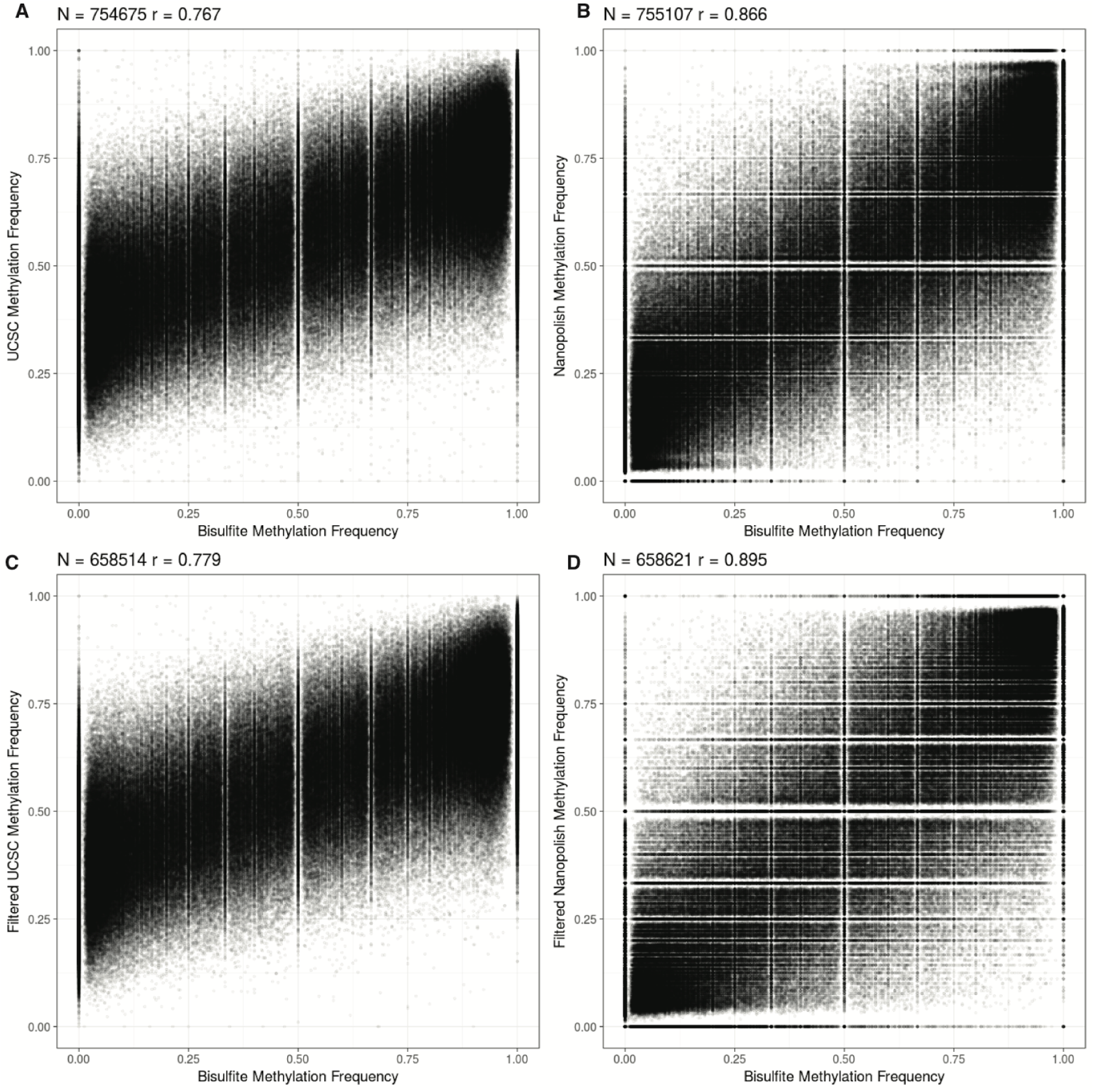
Methylation detection using signal-based methods. A) SignalAlign methylation probabilities compared to bisulfite sequencing frequencies at all called sites. B) Nanopolish methylation frequencies compared to bisulfite sequencing at all called sites. C) SignalAlign methylation probabilities compared to bisulfite sequencing frequencies at sites covered by at least 10 reads in the nanopore and bisulfite data sets, reads were not filtered for quality.

### Ultra-long reads to improve assembly contiguity

Finally, we investigated the impact of read length on the contiguity of our assembly. A 50× PacBio GM12878 dataset with average read length of 4.5 kb previously assembled with an NG50 contig size of 0.9 Mbp ^5^, a third of our assembly. Newer PacBio human assemblies, with mean read lengths greater than 10 kb, have reached contig NG50s exceeding 20 Mbp at 60× coverage ^22^. To project future improvement of nanopore assemblies, we modelled the contribution of read length on assembly, concluding that ultra-long reads would significantly improve assembly continuity (Figure 5A). We therefore developed a method to obtain ultra-long reads by saturating the standard Oxford Nanopore Rapid Kit with high molecular weight DNA (SI Figure 10). We obtained an additional 5× coverage of the genome mainly using this approach (with two additional flowcells employing standard protocols to act as controls because new versions of MinKNOW and an alternative basecaller, Albacore, was used). The N50 read length of the ultraread dataset is 99.7 kb (Figure 5B). The longest full-length mapped read in the dataset (aligned with GraphMap ^31^) is 882kb, corresponding to reference span of 993kb. With the addition of these ultra-long reads, even at low coverage, the nanopore assembly NG50 increased to 6.4 Mbp, more than doubling the previous assembly NG50 and resolving the MHC into a single contig (Figure 5C).

**Figure 5.**
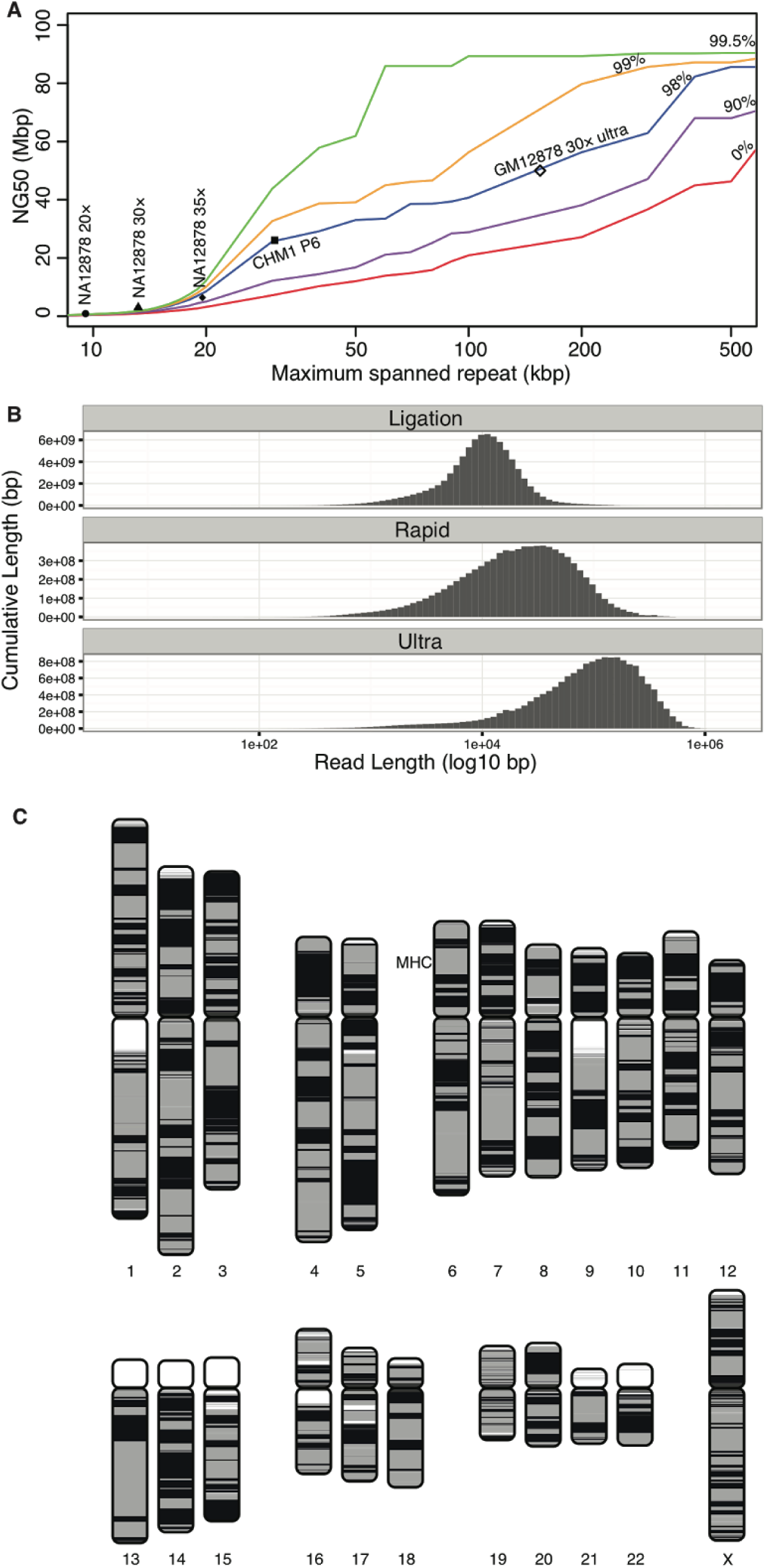
Repeat modeling and assembly. A) A model of expected NG50 contig size when human repeats of a certain length and identity can be correctly resolved (Methods). Repeats can be resolved either by long reads that completely span the repeat, or by accurate reads that can differentiate between non-identical copies. In this simple model, the y-axis shows the expected NG50 contig size when repeats of a certain length (x-axis) or sequence identity (colored lines) can be consistently resolved. Nanopore assembly continuity (GM12878 20×, 30×, 35×) is currently limited by low coverage of long reads and a high error rate, making repeat resolution difficult. These assemblies approximately follow the predicted assembly continuity. The *projected* assembly continuity using 30× of ultra-long reads (GM12878 30× ultra) exceeds 30 Mbp. A recent assembly of 65× PacBio P6 data with an NG50 of 26 Mbp is shown for comparison (CHM1 P6). B) Yield by read length (log10) for ligation, rapid and ultra-long rapid library preparations. The longest reads were achieved via DNA extraction direct from cells using a modified ONT Rapid Sequencing Kit protocol (Methods). C) Chromosomes plot illustrating the continuity of the nanopore assembly boosted with ultra-long reads. Contig NG50 was 6.4 Mbp. Contig and alignment boundaries are represented by a color switch, so regions of continuous color indicate regions of continuous sequence. White areas indicate unmapped sequence, usually caused by N’s in the reference genome. The MHC region on chromosome 6 is labeled, which is contained within a single 15 Mbp contig in the assembly.

### Discussion

We demonstrate the sequencing and assembly of a human genome to high accuracy and contiguity using native DNA and nanopore reads alone. Given approximately 30× coverage the resultant assembly had an identity of 95.74% to the GM12878 reference. Canu was about fourfold slower on the Nanopore data compared to similar coverage of PacBio, requiring ∼60,000 CPU hours for the 30-fold assembly (Methods). This increase is primarily due to systematic error leading to lower accuracy of the corrected reads. While corrected PacBio reads are typically >99% identity, these nanopore reads averaged 92% after correction (SI Figure 1B). Homopolymers represent a challenge for the MinION due to the physical characteristics of the nanopore. Scrappie, developed with this nuance in mind, improves the underlying identity of the chromosome 20 assembly from 95.50% to 97.80%. Consistent with the view that the underlying signal contains additional information, signal based polishing improves the assembly accuracy to 99.44%. Combining signal based polishing and short-read (Illumina) based correction gives a maximum identity of 99.96% or QV34.

We have shown that read lengths on the nanopore platform are closely linked to the input fragment length by generating 5× coverage of ultra-long reads. We demonstrate that careful preparation of DNA in solution using classical extraction and purification methods can yield extremely long read lengths. The longest read lengths were achieved using the transposase based rapid library kit in conjunction with methods of DNA extraction designed to mitigate shearing. This 35× coverage assembly resulted in an NG50 of 6.4 Mb. Based on our modelling observations we predict that 30× of ultra-long reads alone would result in an assembly with a contig NG50 in excess of 40Mb, approaching the continuity of the current human reference, although we have not yet tested this projection (Figure 5C). We speculate that there is no intrinsic read length limit of pore-based systems, other than from physical forces resulting in DNA fragmentation in solution. Therefore there is scope to improve the read length results obtained here further, perhaps through solid phase extraction and library preparation techniques such as use of agar encasement.

Whilst MinION throughput has grown rapidly since its introduction, computational tools to handle and process the data have been slow to scale. We had to develop custom tools to track the large number of reads, each stored as an individual file, and used cloud-based pipelines for much of the analysis (Methods). Free dissemination of these tools as well as their continued engineering support will be invaluable to enable researchers in the future. However, the approach used to sequence a human genome in this study is not likely to represent a convenient method for other users to generate similar data. We anticipate that improvements to the workflow (real-time base-calling, bulk collection files) will be required. A more compact and convenient format for storing raw and base-called data is urgently required, ideally employing a standardised, streaming compatible serialization format such as BAM/CRAM. We were unable to complete an alignment of the ultralong reads using BWA-MEM, suggesting alternative algorithms may be necessary ^32,33^. Additionally, the longest reads exceed CIGAR string limitations in the BAM format, necessitating the use of SAM or CRAM (https://github.com/samtools/hts-specs/issues/40).

The Oxford Nanopore sequencing platform continues to develop at a remarkable pace, making it difficult to make definitive statements on its future potential, and potential ceiling of performance. Here we have demonstrated that sequencing and assembly of a whole human genome using nanopore reads alone is currently possible by running multiple flowcells. We observe that platform throughput continues to improve, with individual flowcells generating >5 Gb of data at best, representing about 12.5% of the theoretical capacity of a 100% efficient flowcell running at 450 bases/second for 48 hours. Higher yields (over 10Gb per flowcell) have been reported by other groups. It remains to be demonstrated how close to full efficiency individual flowcells can be taken consistently through system optimization and adjustment.

Single read accuracy of the MinION technology, in common with other single molecule sequencers, continues to lag behind short-read instruments that achieve better signal discrimination through reading thousands of clonal molecules at a time. Despite this, we are optimistic that consensus accuracy can reach the required Q40 ‘finishing standard’ through platform improvements. We note that template reads have improved significantly since the MinION platform release through a combination of pore changes and bioinformatics improvements. Reads are already sufficiently accurate for highly contiguous *de novo* assembly without complementary Illumina data. The predominant error mode observed in nanopore sequencing are deletions, with particular difficulty around homopolymer sequences. However, the newest base-caller, Scrappie, appears to have made significant progress on this problem by exploiting more of the available nanopore signal. It is reasonable to assume further accuracy gains will be obtained by further exploitation of nanopore signal, including the raw unsegmented electrical current signal data.

Nanopore genotyping accuracy currently lags behind short-read sequencing instruments, particularly due to its limited ability to discriminate between heterozygous and homozygous alleles. We predict genotyping accuracy and variant calling will be improved through better single-read accuracy, increased genome coverage and the utilization of phase constraints between variant sites linked by long reads. Utilization of 1D^2 chemistry that sequences template and complement strands of the same molecule, or better modelling of the nanopore signal data, perhaps incorporating training data from modified DNA, could lead to increased read accuracy. Lastly, increased coverage will be obtained by higher throughput instruments such as GridION and PromethION (respectively equivalent to 5 and ∼130 simultaneous MinION flowcells). Given the high contiguity assemblies we anticipate, structural variant detection and detection of epigenetic marks is a promising early application of this technology, with implications for our understanding of human genetics and, in particular, cancer detection and etiology.

## Methods

### Human DNA input

Human genomic DNA from the GM12878 human cell line (Ceph/Utah pedigree) was either purchased from Coriell (cat no NA12878) or extracted from the cultured cell line. Cell culture was performed using EBV transformed B lymphocyte culture from the GM12878 cell line in RPMI-1640 media with 2mM L-glutamine and 15% fetal bovine serum at 37°C.

### QIAGEN DNA extraction

DNA was extracted from cells using the QIAamp DNA mini kit (Qiagen). 5×10^6^ cells were spun at 300× *g* for 5 minutes to pellet. The cells were resuspended in 200 μl PBS and DNA was extracted according to the manufacturer’s instructions. DNA quality was assessed by running 1 μl on a genomic ScreenTape on the TapeStation 2200 (Agilent) to ensure a DNA Integrity Number (DIN) >7 (Value for NA12878 was 9.3). Concentration of DNA was assessed using the dsDNA HS assay on a Qubit fluorometer (Thermo Fisher).

### Library preparation (SQK-LSK108 1D ligation genomic DNA)

1.5–2.5 μg human genomic DNA was sheared in a Covaris g-TUBE centrifuged at 5000–6000 rpm in an Eppendorf 5424 (or equivalent) centrifuge for 2× 1 minute, inverting the tube between centrifugation steps.

DNA repair (NEBNext FFPE DNA Repair Mix, NEB M6630) was performed on purchased DNA but not on freshly extracted DNA. 8.5 μl NFW, 6.5 μl FFPE Repair Buffer and 2 μl FFPE DNA Repair Mix were added to the 46 μl sheared DNA. The mixture was incubated for 15 mins at 20 °C, cleaned up using a 0.4× volume of AMPure XP beads (62 μl), incubated at room temperature with gentle mixing for 5 minutes, washed twice with 200 μl fresh 70% ethanol, pellet allowed to dry for 2 mins and DNA eluted in 46 μl NFW or EB (10 mM Tris pH 8.0). A 1 μl aliquot was quantified by fluorometry (Qubit) to ensure ≥1 μg DNA was retained.

End repair and dA-tailing (NEBNext Ultra II End-Repair / dA-tailing Module) was then performed by adding 7 μl Ultra II End-Prep buffer, 3 μl Ultra II End-Prep enzyme mix, and 5 μl NFW. The mixture was incubated at 20 °C for 10 minutes and 65 °C for 10 minutes. A 1× volume (60 μl) AMPure XP clean-up was performed and the DNA was eluted in 31 μl NFW. A 1 μl aliquot was quantified by fluorometry (Qubit) to ensure ≥700 ng DNA was retained.

Ligation was then performed by adding 20 μl Adapter Mix (SQK-LSK108 Ligation Sequencing Kit 1D, Oxford Nanopore Technologies [ONT]) and 50 μl NEB Blunt/TA Master Mix (NEB, cat no M0367) to the 30 μl dA-tailed DNA, mixing gently and incubating at room temperature for 10 minutes.

The adapter-ligated DNA was cleaned-up by adding a 0.4× volume (40 μl) of AMPure XP beads, incubating for 5 minutes at room temperature and resuspending the pellet twice in 140 μl ABB (SQK-LSK108). The purified-ligated DNA was resuspend by adding 25 μl ELB (SQK-LSK108) and resuspending the beads, incubating at room temperature for 10 minutes, pelleting the beads again and transferring the supernatant (pre-sequencing mix or PSM) to a new tube. A 1 μl aliquot was quantified by fluorometry (Qubit) to ensure ≥ 500 ng DNA was retained.

### Sambrook and Russell DNA extraction

This protocol was modified from Chapter 6 protocol 1 of Sambrook and Russell ^34^. 5×10^7^ cells were spun at 4500× *g* for 10 minutes to pellet. The cells were resuspended by pipette mixing in 100 μl PBS. 10ml TLB was added (10mM Tris-Cl pH 8.0, 25mM EDTA pH 8.0, 0.5% (w/v) SDS, 20 μg/ml Qiagen RNase A), vortexed at full speed for 5 seconds and incubated at 37 °C for 1 hr. 50 μl Proteinase K (Qiagen) was added and mixed by slow inversion 10 times followed by 3 hrs at 50 °C with gentle mixing every 1 hour. The lysate was phenol purified using 10 ml buffer saturated phenol using phase-lock gel falcon tubes, followed by phenol:chloroform (1:1), The DNA was precipitated by the addition of 4 ml 5 M ammonium acetate and 30 ml ice-cold ethanol. DNA was recovered with a glass hook followed by washing twice in 70% ethanol. After spinning down at 10,000g, ethanol was removed followed by 10 mins drying at 40 °C. 150 μl EB was added to the DNA and left at 4 °C overnight to resuspend.

### Library preparation (SQK-RAD002 genomic DNA)

To obtain ultra-long reads, the standard RAD002 protocol (SQK-RAD002 Rapid Sequencing Kit, ONT) for genomic DNA was modified as follows. 16 μl of DNA from the Sambrook extraction at approximately 1 μg/μl, manipulated with a cut-off P20 pipette tip, was placed in a 0.2 ml PCR tube, with 1 μl removed to confirm quantification value. 5 μl FRM was added and mixed slowly 10 times by gentle pipetting with a cut-off pipette tip moving only 12 μl. After mixing, the sample was incubated at 30 °C for 1 minute followed by 75 °C for 1 minute on a thermocycler. After this, 1 μl RAD and 1 μl Blunt/TA ligase was added with slow mixing by pipetting using a cut-off tip moving only 14 μl 10 times. The library was then incubated at room temperature for 30 minutes to allow ligation of Rapid Adapters (RAD). To load the library, 25.5 μl RBF was mixed with 27.5 μl NFW and this was added to the library. Using a P100 cut-off tip set to 75 μl, this library was mixed by pipetting slowly 5 times. This extremely viscous sample was loaded onto the “spot on” port and entered the flow cell by capillary action. The standard loading beads were omitted from this protocol due to excessive clumping when mixed with the viscous library

### MinION sequencing

MinION sequencing was performed as per manufacturer’s guidelines using R9/R9.4 flowcells (FLO-MIN105/FLO-MIN106, ONT). Reads from all sites were copied off to a volume mounted on a CLIMB virtual server (http://www.climb.ac.uk) where metadata was extracted using poredb (https://github.com/nickloman/poredb) and base-calling performed using Metrichor (predominantly workflow ID 1200 although previous versions were used early on in the project), Nanonet (https://github.com/nanoporetech/nanonet) and Scrappie (ONT) were used for the chr20 comparisons using reads previously identified as from this chromosome after mapping the Metrichor reads. Albacore 0.8.4 (available from the Oxford Nanopore Technologies user community) was used for the ultralong read set, as this software became the recommended basecaller for nanopore reads in March 2017.

### Modified MinION running scripts

In a number of instances, MinION sequencing control was shifted to non-standard MinKNOW scripts. These scripts provided enhanced pore utilisation/data yields during sequencing, and operated by monitoring and adjusting flowcell bias-voltage, active pore reselection, and event yield dependent adjustment while still active. More detailed information on these scripts can be found on the Oxford Nanopore Technologies user community.

### Live run monitoring

To assist in choosing when to switch from a standard run script to a modified run protocol, a subset of runs were monitored with the assistance of the minControl tool, an alpha component of the minoTour suite of minION run and analysis tools (https://github.com/minoTour/minoTour). minControl collects metrics about a run directly from the grouper software, which runs behind the standard ONT MinKNOW interface. minControl provides a historical log of yield measured in events from a flowcell enabling estimations of yield and the decay rate associated with loss of sequencing pores over time. MinKNOW yield is currently measured in events and is scaled by approximately 1.7 to estimate yield in bases.

### Assembly

All “NG” statistics were computed using a genome size of 3,098,794,149 bp (3.1 Gbp), the size of GRCh38 excluding alt sites.

Canu v1.4 (+11 commits) r8006 (4a7090bd17c914f5c21bacbebf4add163e492d54) was used to assemble the initial 20-fold coverage dataset:

~~~
canu -p asm -d asm genomeSize=3.1g gridOptionsJobName=na12878nano “gridOptions=--time 72:00:00 --partition norm” -nanopore-raw rel2*.fastq.gz corMinCoverage=0 corMaxEvidenceErate=0.22 errorRate=0.045
~~~

These are the suggested low-coverage parameters from the Canu documentation, but with a decreased maximum evidence error rate. This specific parameter was lowered to reduced memory requirements after it was determined that the MinHash overlapping algorithm was under-estimating error rates due to systematic error in the reads. Counterintuitively, this systematic error makes two reads look more similar than reality because they share more *k*-mers than expected under a random model. Manually decreasing the maximum overlap error rate threshold adjusted for this bias. The assembly took 40K CPU hours (25K to correct and 15K to assemble). This is about twofold slower than a comparable PacBio dataset, mostly due to the higher noise and systematic error in the nanopore reads.

The same version of Canu was also used to assemble the 30-fold dataset:

~~~
canu -p asm -d asm genomeSize=3.1g gridOptionsJobName=na12878nano “gridOptions=--time 72:00:00 --partition norm” -nanopore-raw rel3*.fastq.gz corMinCoverage=0 corMaxEvidenceErate=0.22 errorRate=0.045 “corMhapOptions=--threshold 0.8 --num-hashes 512 --ordered-sketch-size 1000 --ordered-kmer-size 14”
~~~

For this larger dataset, overlapping was again tweaked by reducing the number of hashes used and increasing the minimum overlap identity threshold. This has the effect of lowering sensitivity to further compensate for the bias in the input reads. This assembly required 62K CPU hours (29K to correct, 33K to assemble), which is about fourfold slower than a comparable PacBio dataset.

The combined dataset incorporating an additional 5× coverage of ultra-long reads was assembled with an updated version of Canu v1.4 (+125 commits) r8120:

~~~
canu -p asm -d asm genomeSize=3.1g gridOptionsJobName=na12878nano “gridOptions=--time 72:00:00 --partition norm” -nanopore-raw rel3*.fastq.gz -nanopore-raw rel4*.fastq.gz
“corMhapOptions=--threshold 0.8 --num-hashes 512
--ordered-sketch-size 1000 --ordered-kmer-size 14” batOptions=“-dg
3 -db 3 -dr 1 -el 2000 -nofilter suspicious-lopsided”
~~~

This assembly required 151K CPU hours (15K to correct, 86K to trim, and 50K to assemble). These high runtimes are a consequence of the ultra-long reads. In particular, the current Canu trimming algorithm was not designed for reads of this extreme length and high error rate and the algorithms used are not optimal. The future performance of nanopore assembly will undoubtedly improve with tailored algorithms and improved base calling accuracy.

### Assembly continuity modeling

Expected assembly continuity was modeled on repeat tracks downloaded from the UCSC genome browser (http://hgdownload.soe.ucsc.edu/goldenPath/hg38/database/).

For a given repeat identity (0%, 90%, 95%, 98%, 99%, and 99.5%), all repeats with a lower identity estimate (genomicSuperDups and chainSelf) were filtered and overlapping repeats were merged. Gaps in the reference were also considered as repeats. To compute the maximum repeat length likely to be spanned by a given sequence distribution, the probability of an unspanned repeat of a fixed length was estimated for all lengths between 1 and 100 kbp in steps of 1 kbp using an equation from http://data-science-sequencing.github.io/lectures/lecture7/ ^35–37^:

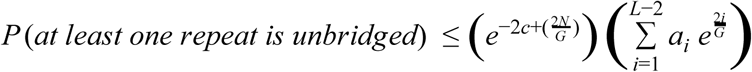

where *G* is the genome size, *L* is the read length, *ai* is the number of repeats of length 1 ≤ *i* ≤ *L* − 2, *N* is the number of reads ≥ *L*, and *c* is the coverage in reads ≥ *L*. We used the distribution of all repeats for *a_i_* and plotted the shortest repeat length such that *P* (*at least one repeat is unbridged*) > 0.05 for real sequencing length distributions both nanopore and PacBio sequencing runs. Assemblies of the data were plotted at their predicted spanned read length on the x-axis and NG50 on the y-axis for comparison with the model. A 30× run of ultra-long coverage was simulated from the 5× dataset by repeating each ultra-long read six times.

### Assembly validation and structural variant analysis

Assemblies were aligned using MUMmer v3.23 with parameters “-l 20 -c 500 -maxmatch” for the raw assemblies and “-l 100 -c 500 -maxmatch” for the polished assemblies. Output was processed with dnadiff to report average 1-to-1 alignment identity. The MUMmer coords file was converted to a tiling using the scripts from Berlin *et al.* ^38^ with the command:

~~~
python convertToTiling.py 10000 90 100000
~~~

and drawn using the coloredChromosomes package ^39^. Since the reference is a composite of human genomes and there are true variations between the reference and NA12878, we also computed a reference-free estimate of identity. A 30-fold subset of the Genome In a Bottle Illumina dataset for NA12878 ^17^ was downloaded from ftp://ftp-trace.ncbi.nlm.nih.gov/giab/ftp/data/NA12878/NIST_NA12878_HG001_HiSeq_300x/RMNISTHS_30xdownsample.bam. Samtools fastq was used to extract fastq paired-end data for the full dataset and for the reads mapping to chromosome 20.

The reads were aligned to the whole genome assembly and chromosome 20 assemblies with BWA-MEM 0.7.12-r1039. Variants were identified using FreeBayes v1.0.2 ^40^ with the command:

~~~
freebayes -C 2 -0 -O -q 20 -z 0.10 -E 0 -X -u -p 2 -F 0.6 -b alignments.bam -v asm.bayes.vcf -f asm.fasta
~~~

The length of all variants was summed and the total number of bases with at least 3× coverage was summed using samtools depth. QV was computed as 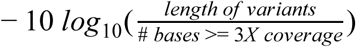, and identity was computed as 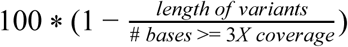. Dotplots were generated with “mummerplot --fat” using the 1-to-1 filtered matches.

A previously published GM12878 PacBio assembly ^5^ was aligned as above with MUMmer v3.23. The resulting alignment files were uploaded to Assemblytics ^41^ to identify structural variants and generate summary figures. Versus GRCh38, the PacBio assembly identified 10,747 structural variants affecting 10.84 Mbp, and reported an equal balance of insertions and deletions (2,361 vs. 2,724), with a peak at approximately 300 bp corresponding to Alu repeats (SI Figure 5 A, SI Table 6). The high error rate of the nanopore assembly resulted in a much larger number of identified variants (69,151) affecting 23.45 Mbp, with a strong deletion bias (3,900 insertions vs. 28,791 deletions) (SI Figure 5B, SI Table 6). The Illumina-polished assembly reduced the total variants (47,073) affecting 16.24 Mbp but the deletion bias persisted (2,840 insertions vs. 20,797 deletions) (SI Figure 5 C, SI Table 6).

### Base call analysis

Sequences were aligned to the 1000 genome GRCh38 reference using BWA-MEM version 0.7.12-r1039 with the “-x ont2d” option ^42^. The BAM alignments were converted to PAF format ^32^ and cigar-strings parsed to convert alignments to an identity. Summary statistics for each flowcell were tabulated separately and combined. Alignment length versus identity was plotted using smoothScatter in R. Depth of coverage statistics for each flowcell were obtained from “samtools depth -a” and combined. As for the assembly statistics, a genome size of 3,098,794,149 bp was used to compute bases covered. The mean coverage was 25.63 (63.20 sd). The minimum coverage was 0 and the maximum was 44,391. Excluding 0-coverage regions, the mean coverage was 27.41 (64.98 sd). The coverage histogram was plotted compared with randomly-generated Poisson values generated with R’s rpois function with λ = 27.4074.

Metrichor reads mapping to human chromosome 20 were additionally base-called with Nanonet v2.0 and Scrappie v0.2.7. Scrappie reads comprised primarily of low-complexity sequence were identified using the sdust program included with Minimap (commit: 17d5bd12290e0e8a48a5df5afaeaef4d171aa133) ^32^ with default parameters (-w 64 -t 20). The total length of the windows in a single sequence were merged and divided by read length to compute percentage of low-complexity sequence in each read. Any read for which this percentage exceeded 50% was removed from downstream analysis. Without this filtering, BWA-MEM did not complete mapping the sequences after >30 days of runtime on 16-cores.

To measure homopolymer accuracy, pairwise read-to-reference alignments were extracted for reads spanning all homopolymers of length 2 or greater. For efficiency, at most 1000 randomly selected instances were considered for each homopolymer length. Each homopolymer so-identified is enclosed by two non-homopolymer “boundary” bases (for example, the T and G in TAAAG). The number of match, mismatch, insertion and deletion alignment operations between the boundary bases was tabulated for each homopolymer, and alignments not anchored at the boundary bases with match/mismatch operations were ignored. Homopolymer call length was reported as the number of inserted bases minus the number of deleted bases in the extracted alignment, quantifying the difference between expected and observed sequence length. All base callers with the exception of Scrappie failed in large homopolymer stretches (e.g. SI Figure 3), consistently capping homopolymers at 5 bp (the *k*-mer length of the model). Scrappie shows significant improvement, but tended to slightly over-call short homopolymers and under-call longer ones (Figure 2B).

To quantify deviations from the expected 50/50 allele ratio at heterozygous sites, 25,541 homozygous and 46,098 heterozygous SNP positions on chromosome 20 were extracted from the Illumina Platinum Genomes project VCF for GM12878, requiring a minimum distance of 10 bp between SNP positions. Scrappie base calls at these positions were extracted using samtools mpileup. Deviation from the expected allelic ratio was defined as *d* = abs(0.5 - [allele A coverage]/[allele A coverage + allele B coverage]). Averaged over all evaluated heterozygous SNPs, *d* = 0.13 and 90% of SNPs have *d* <= 0.27 (corresponding to approximately >= 25% coverage on the minor allele). Results were similar when stratified by SNP type.

### Assembly polishing with Nanopolish

We ran the nanopolish consensus calling algorithm on the three chromosome 20 assemblies described above. For each assembly we sampled candidate variants from the base-called reads used to construct the contigs (using the “--alternative-basecalls” option) and input the original fast5 files (generated by the basecaller in the Metrichor computing platform) into a hidden Markov model, as these files contained the annotated events that the HMM relies on. The reads were mapped to the draft assembly using BWA-MEM with the “-x ont2d” option.

Each assembly was polished in 50,000 bp segments and the individual segments were merged into the final consensus. The nanopolish jobs were run using default parameters except the “--fix-homopolymers” and “--min-candidate-frequency 0.01” options were applied.

### Assembly annotation

Comparative Annotation Toolkit (CAT) (https://github.com/ComparativeGenomicsToolkit/Comparative-Annotation-Toolkit commit c9503e7) was run on both the polished and unpolished assemblies. CAT uses whole genome alignments to project transcripts from a high-quality reference genome to other genomes in the alignment ^43^. The gene finding tool AUGUSTUS is used to clean up these transcript projections and a combined gene set is generated^44^.

To guide the annotation process, human RNA-seq data were obtained from SRA for a variety of tissues and aligned to both hg38 and the two assembly versions. GENCODE V24 was used as the reference annotation. Two separate progressiveCactus ^45^ alignments were generated for each assembly version with the chimpanzee genome as an outgroup.

The frequency of frameshifting insertions or deletions (Indels) in transcripts was evaluated by performing pairwise CDS sequence alignments using BLAT in a codon-aware parameterization. Alignments were performed both on raw transMap output as well as on the final consensus transcripts. The observed rate of coding insertions compared to deletions was not equal -- 29% of transMap transcripts had a frameshifting insertion, and 50% had a frameshifting deletion, suggesting a systematic over-representation of spurious deletions.

### MHC analysis

Exon sequences belonging to the six classical HLA genes were extracted from the Illumina-polished assembly, and HLA types called at G group resolution. These results were compared to GM12878 HLA type reference data. For the class I HLA genes (represented with one copy each), there was good agreement between the best-matching reference type and the alleles called from the assembly (edit distance 0–2). For the class II HLA genes, there was perfect agreement for 2 HLA-DQA1 and the 2 HLA-DQB1 alleles, providing further evidence for the presence of a correctly assembled class II haplotype fragment in the assembly. Detailed examination of HLA-DRB1, however, showed that one exon is largely absent from the assembly. The presence of a deletion or a non-canonical haplotype structure around the HLA-DRB homologs in GM12878 is consistent with Dilthey *et al.* in which a reference-based approach also failed to resolve the structure of the class II region near the HLA-DRB homologs ^46^.

For HLA typing, contigs from the MHC region were identified using MUMmer and then globally aligned to each of 8 MHC ALT haplotypes in GRCh38. Global alignments were computed using BWA-MEM seed alignments with parameters “-a -x pacbio”, followed by dynamic programming to identify an optimal alignment, restricting the set of considered Needleman-Wunsch paths to the seed diagonals and connections between them. For each contig, the ALT haplotype with the best alignment score was selected and annotations projected onto it using the gene annotation set underlying the HLA*PRG graph ^47^. To carry out HLA typing from a contig for a specific gene at G group resolution, contigs were required to contain the relevant exons (exons 2 and 3 for class I HLA genes, and exon 2 for class II HLA genes). Exon sequences were extracted and the G group ^48^ exhibiting minimum edit distance to the extracted sequences was reported. For one contig with hits to both HLA-DRB1 and HLA-DRB3, MAFFT ^49^ (with --auto) was used to validate and refine the alignment. GM12878 G group HLA types for HLA-A, -B, -C, -DQA1, -DQB1 and -DRB1 are from ^46^; the presence of exactly one HLA-DRB3 allele is expected due to linkage with HLA-DRB1 (DRB1*03 is associated with HLA-DRB3, and DRB1*01 has no DRB3/4/5 association), and was confirmed with HLA*PRG (SI Table 7).

To phase the large contig (tig00019339) spanning the MHC class II region, heterozygous sites were extracted by mapping Illumina reads to the polished assembly using BWA-MEM with default parameters. Alignments were post-processed according to the GATK 3.7 whole-genome variant calling pipeline, except for the “-T IndelRealigner” step using “--consensusDeterminationModel USE_READS”. The -T HaplotypeCaller parameter was used for variant calling. Nanopore reads were aligned back to the assembly using BLASR ^50^ and the combined VCF file used for phasing. WhatsHap ^51^ with the “-indels” option was used to extract phasing marking variants, and reads with more than 1 phasing marking were classified as haplotype A or B when >65% of their variants were in agreement (SI Figure 6).

### Genotyping SNPs using Nanopolish

Nanopolish was used for genotyping the subset of reads that mapped to human chromosome 20. The 1000 Genomes phase 3 variant set for GRCh38 was used as a reference and filtered to include only chromosome 20 SNPs that were not singletons (AC ≥ 2). This set of SNPs was input into “nanopolish variants” in genotyping mode (“--genotype”). The genotyping method extends the variant calling framework previously described ^9^ to consider pairs of haplotypes, allowing it to be applied to diploid genomes (option “--ploidy 2”). To evaluate their accuracy, genotype calls were compared to the “platinum calls” generated by Illumina ^20^. When evaluating the correctness of a nanopore call, we required the log-likelihood ratio of a variable call (heterozygous or homozygous non-reference) to be at least 30, otherwise we considered the site to be homozygous reference.

### Estimating SV genotyping sensitivity

2,435 previously identified high-confidence GM12878 SVs marked as “duplications” or “deletions” were used to determine genotype sensitivity ^25^. These “gold standard” SVs were genotyped using SVTyper ^24^ in the Platinum Genomes NA12878 Illumina dataset (paired-end reads; European Nucleotide Archive, Run Accession ERR194147). Nanopore reads were mapped using BWA-MEM and the “-x ont2d” flag resulting in a BAM file for each flowcell. Random subsets of flowcells were then merged to simulate using *X* flowcells worth of data for a given analysis. Gold standard SVs were then genotyped in each merged BAM file using a modified version of SVTyper (http://github.com/tomsasani/svtyper). Generally, long nanopore reads are subject to higher rates of mismatches, insertions, and deletions than short Illumina reads. These features can result in “bleed-through” alignments, where reads align past the true breakpoint of an SV ^52^. The modifications to SVTyper attempt to correct for the “bleed-through” phenomenon by allowing reads to align past the breakpoint, yet still support an alternate genotype. All modifications to SVTyper are documented in the source code available at the GitHub repository listed above. Random BAM merging and genotyping was repeated three times. Nanopore and Illumina derived genotypes were then compared as a function of the number of flowcells.

Nanopore false-discovery rate was estimated by randomly permuting the genomic locations of the original SVs using BEDTools “shuffle” ^53^. Centromeric, telomeric, and “gap” regions (as defined by the UCSC Genome Browser) were excluded when assigning randomly selected breakpoints to each SV. The randomly shuffled SVs were then genotyped in Illumina and nanopore data in the same manner as before. It is expected that the alignments at shuffled SV intervals would almost always support a homozygous reference genotype. So, all instances in which Illumina data supported a homozygous reference genotype, yet the nanopore data called a non-homozygous reference genotype, were considered false positives. SV coordinates were shuffled and genotyped 1000 times and the average false discovery rate over all iterations was 6.4%.

### Scaling marginAlign and signalAlign data analysis pipelines

To handle the large data volume, the original marginAlign and signalAlign algorithms were ported to cloud infrastructures using the Toil batch system ^54^. Toil allows for computational resources to be scaled horizontally and vertically as a given experiment requires and enables researchers to perform their own experiments in identical conditions. All of the workflows used and the source code is freely available from https://github.com/ArtRand/toil-signalAlign and https://github.com/ArtRand/toil-marginAlign. Workflow diagrams are shown in SI Figure 11.

### Generating a controlled set of methylated control DNA samples

DNA methylation control standards were obtained from Zymo Research (cat. Number D5013). The standards contain a whole-genome-amplified (WGA) DNA substrate that lacks methylation and a WGA DNA substrate that has been enzymatically treated so all CpG dinucleotides contain 5-methyl cytosines. The two substrates were sequenced independently in two different flowcells using the sequencing protocol described above. Training for signalAlign and nanopolish was carried out as previously described ^29,30^.

### 5-methyl cytosine detection with signalAlign

The signalAlign algorithm uses a variable order hidden Markov model combined with a hierarchical Dirichlet process (HMM-HDP) to infer base modifications in a reference sequence using the ionic current signal produced by nanopore sequencing^55^. The ionic current signal is simultaneously influenced by multiple nucleotides as the strand passes through the nanopore. Correspondingly, signalAlign models each ionic current state as a nucleotide *k*-mer. The model allows a base in the reference sequence to have any of multiple methylation states (in this case 5-methy cytosine or canonical cytosine). The model ties the probabilities of consistently methylated *k*-mers by configuring the HMM in a variable order meta-structure that allows for multiple paths over a reference *k*-mer depending on the number of methylation possibilities. To learn the ionic current distributions for methylated *k*-mers, signalAlign estimates the posterior mean density for each *k*-mer’s distribution of ionic currents using a Markov chain Monte Carlo (MCMC) algorithm given a set of *k*-mer-to-ionic current assignments. Using the full model, the posterior for each methylation status is calculated for all cytosines in CpG dinucleotides.

### 5-methyl cytosine detection with nanopolish

Previous work describes using nanopolish to call 5-methylcytosine in a CpG context using a hidden Markov model ^30^. The output of the nanopolish calling procedure is a log-likelihood ratio, where a positive log-likelihood ratio indicates evidence for methylation. Nanopolish groups nearby CpG sites together and calls the group jointly, assigning the same methylation status to each site in the group. To allow comparison to the bisulfite data each such group was broken up into its constituent CpG sites, which all have the same methylation frequency. Percent-methylation was calculated by converting the log-likelihood ratio to a binary methylated/unmethylated call for each read, and calculating the fraction of reads classified as methylated. A filtered score was also computed by first filtering reads where the absolute value of the log-likelihood ratio was less than 2.5 to remove ambiguous reads.

## Acknowledgements

We would like to acknowledge the support of Oxford Nanopore Technologies in generating this dataset, with particular thanks to Rosemary Dokos, Oliver Hartwell, Jonathan Pugh, and Clive Brown. We also thank Mark Akeson for his support and insight. We would like to thank Radoslaw Poplawski and Simon Thompson for technical assistance with configuring and using cloud-based file systems with millions of files on CLIMB. We thank Winston Timp and Rachael Workman for generating the R9.4 methylation training data for nanopolish. We thank Thorsten Allers for assistance with PFGE. We would like to thank Angel Pizarro at Amazon Web Services for hosting the human genome dataset as an Amazon Web Services Open Dataset. This study utilized the computational resources of the Biowulf system at the National Institutes of Health, Bethesda, MD (https://biowulf.nih.gov).

## Conflict of interest statement

ML, NL, JOG, JTS, JRT, and TPS were members of the MinION access program (MAP) and have received free-of-charge flowcells and kits for nanopore sequencing for this and other studies, and travel and accommodation expenses to speak at Oxford Nanopore Technologies conferences. NJL has received an honorarium to speak at an Oxford Nanopore company meeting. SK, ATD, and TAS have received travel and accommodation expenses to speak at Oxford Nanopore Technologies conferences.

JTS, JOG and ML receive research funding from Oxford Nanopore Technologies.

## Funding

JOG is supported by the UK Antimicrobial Resistance Cross Council Initiative supported by the seven research councils (MR/N013956/1) and Rosetrees Trust grant A749. SK, AD, AR, and AMP were supported by the Intramural Research Program of the National Human Genome Research Institute, National Institutes of Health. ML is supported by the BBSRC (BB/N017099/1 and BB/M020061/1). JRT and TPS are supported by the Canadian Institutes of Health Research (grant #10677) and a Brain Canada Multi-Investigator Research Initiative Grant with matching support from Genome British Columbia, the Michael Smith Foundation for Health Research and the Koerner Foundation. TPS is also supported by the Canada Research Chair in Biotechnology and Genomics-Neurobiology. JTS is supported by the Ontario Institute for Cancer Research through funds provided by the Government of Ontario. ARQ, BSP, and TAS are supported by a US National Human Genome Research Institute award (NIH R01HG006693) and a US National Cancer Institute award (NIH U24CA209999). ADB acknowledges funding from the Wellcome Trust (102732/Z/13/Z), Cancer Research UK (A23923) and the Medical Research Council UK (MR/M016587/1).

## Supplementary Information

### Supplementary Figures

**SI Figure 1.**
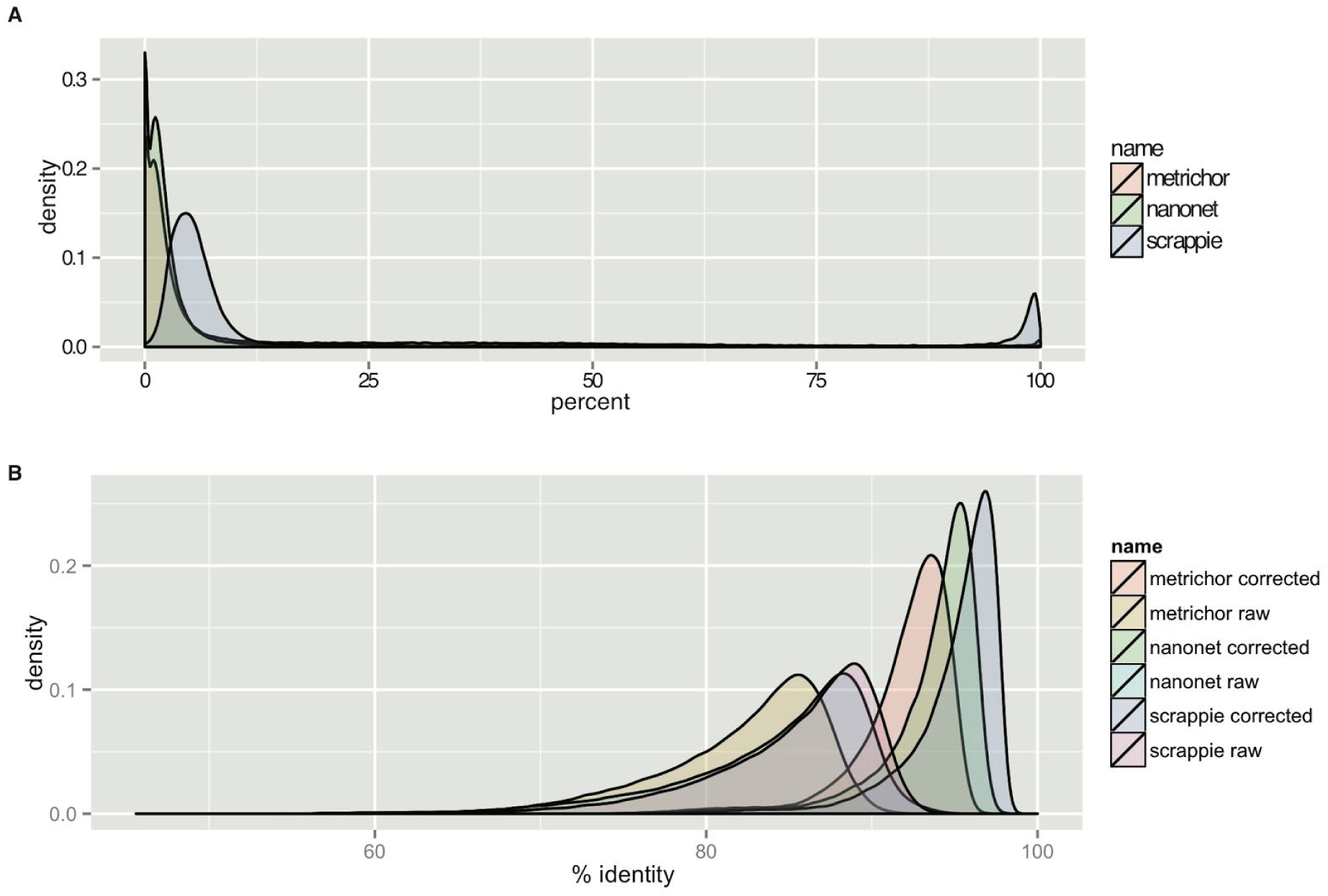
Read Complexity. A) Density plot showing the percentage of read length masked by the ‘dust’ program, which identifies low-complexity sequence (simple repeats). Scrappie outputs a significantly larger fraction of low-complexity bases, including some reads that are entirely low-complexity sequence. B) Density plot showing the % identity for reads, weighted by alignment length, basecalled with metrichor, nanonet and scrappie both pre and post correction.

**SI Figure 2.**
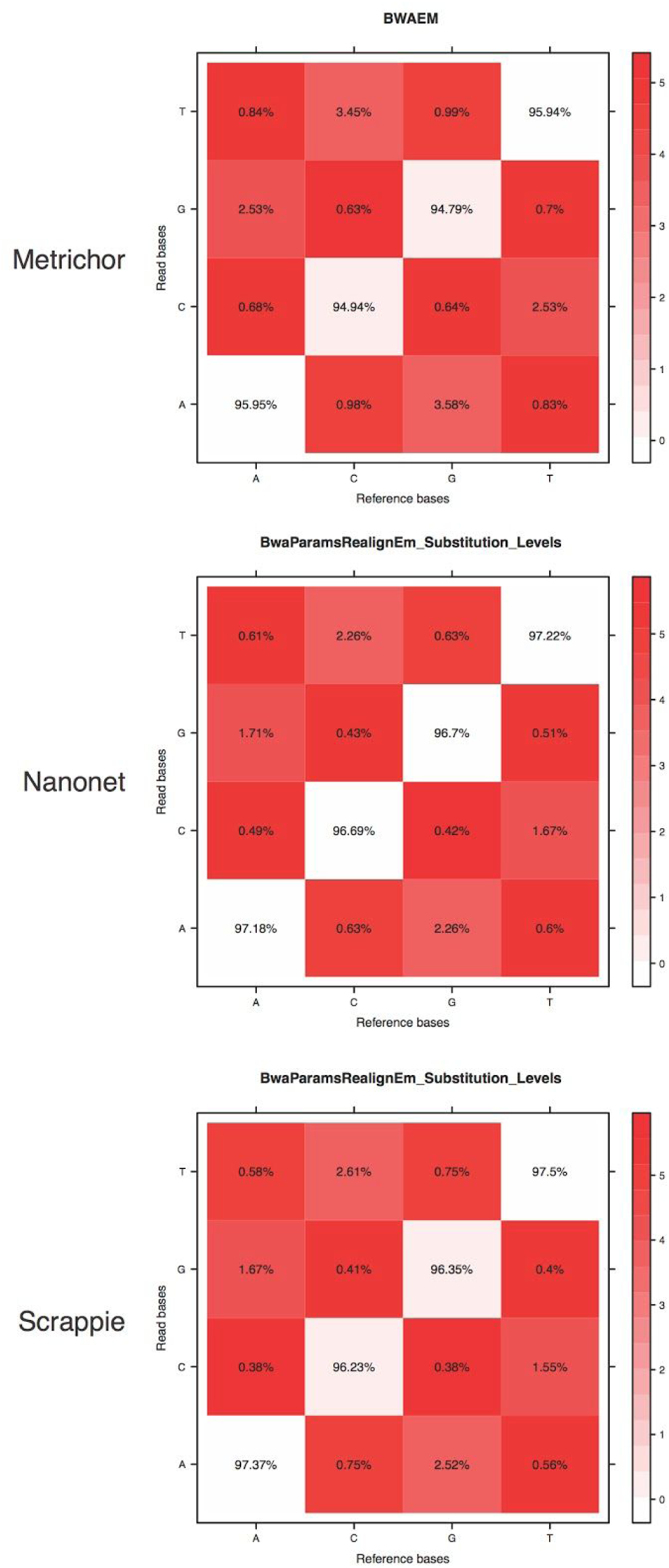
Basecall Bias. Confusion matrices describing call bias for the three base calling algorithms used from high-confidence alignments.

**SI Figure 3.**
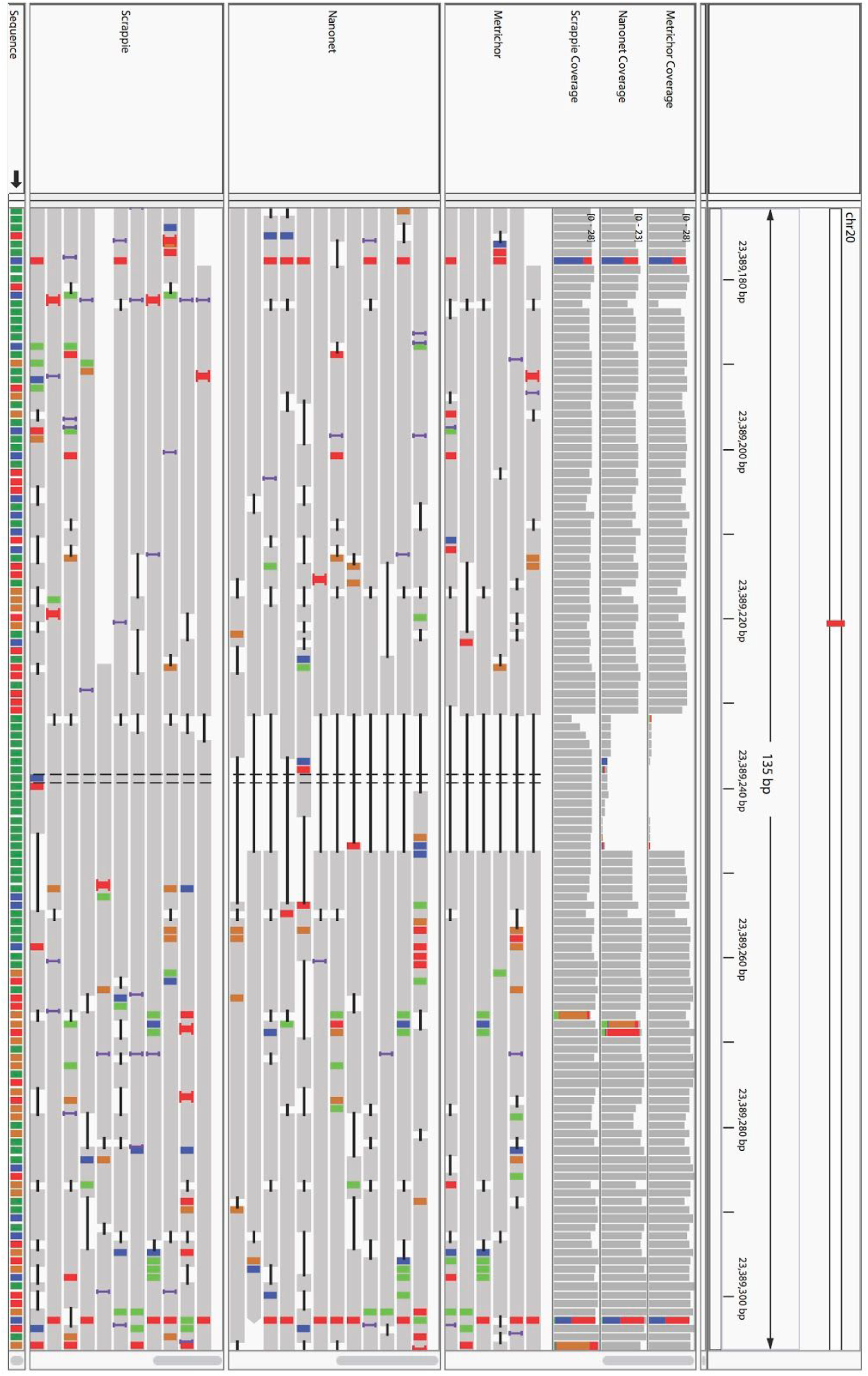
Illustrative homopolymer resolution by basecaller. IGV plot showing a poly-A region and aligned reads from Metrichor, Nanonet, and Scrappie base callers. The top three tracks show coverage across the region, and the bottom three tracks show the read alignments. Horizontal black bars in the read alignment tracks indicate deletions. Colorful bars indicate mismatches. Metrichor and Nanonet fail to call the homopolymer entirely, but Scrappie produces more reasonable calls across this region.

**SI Figure 4.**
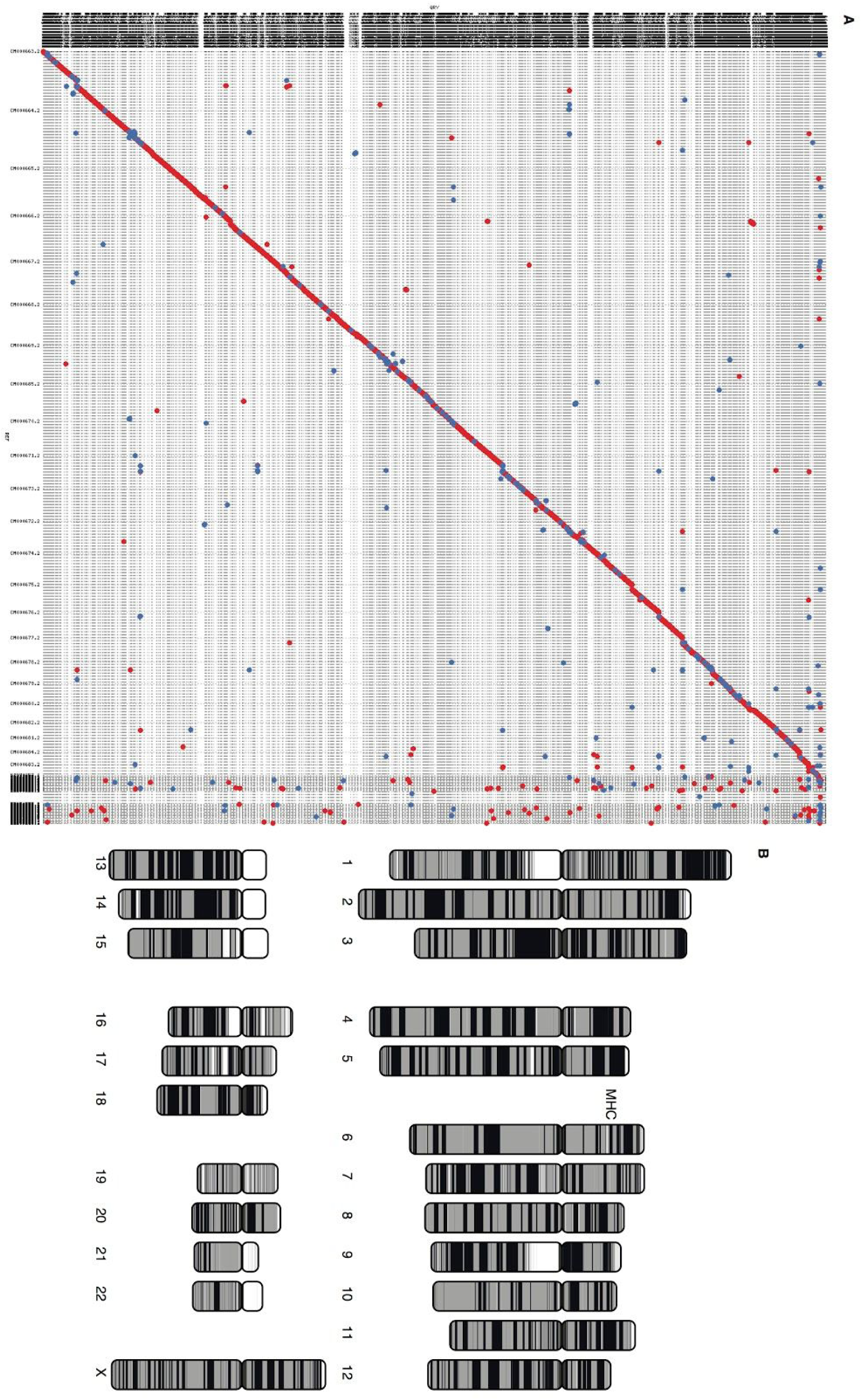
Assembled contigs against reference. A) Alignment dotplot of the nanopore GM12878 assembly aligned against human reference GRCh38 showing overall structural agreement. Human chromosomes are arranged along the x-axis with assembled contigs along the y-axis. Grid lines indicate chromosome and contig boundaries. Forward-strand matches are in red and reverse-complement in blue. B) Chromosomes plot illustrating the continuity of the 30× nanopore assembly. Contig NG50 was 3 Mbp. Contig and alignment boundaries are represented by a color switch, so regions of continuous color indicate regions of continuous sequence. White areas indicate unmapped sequence, usually caused by N’s in the reference genome. The MHC region on chromosome 6 is labeled, which is reconstructed as described in the main text.

**SI Figure 5.**
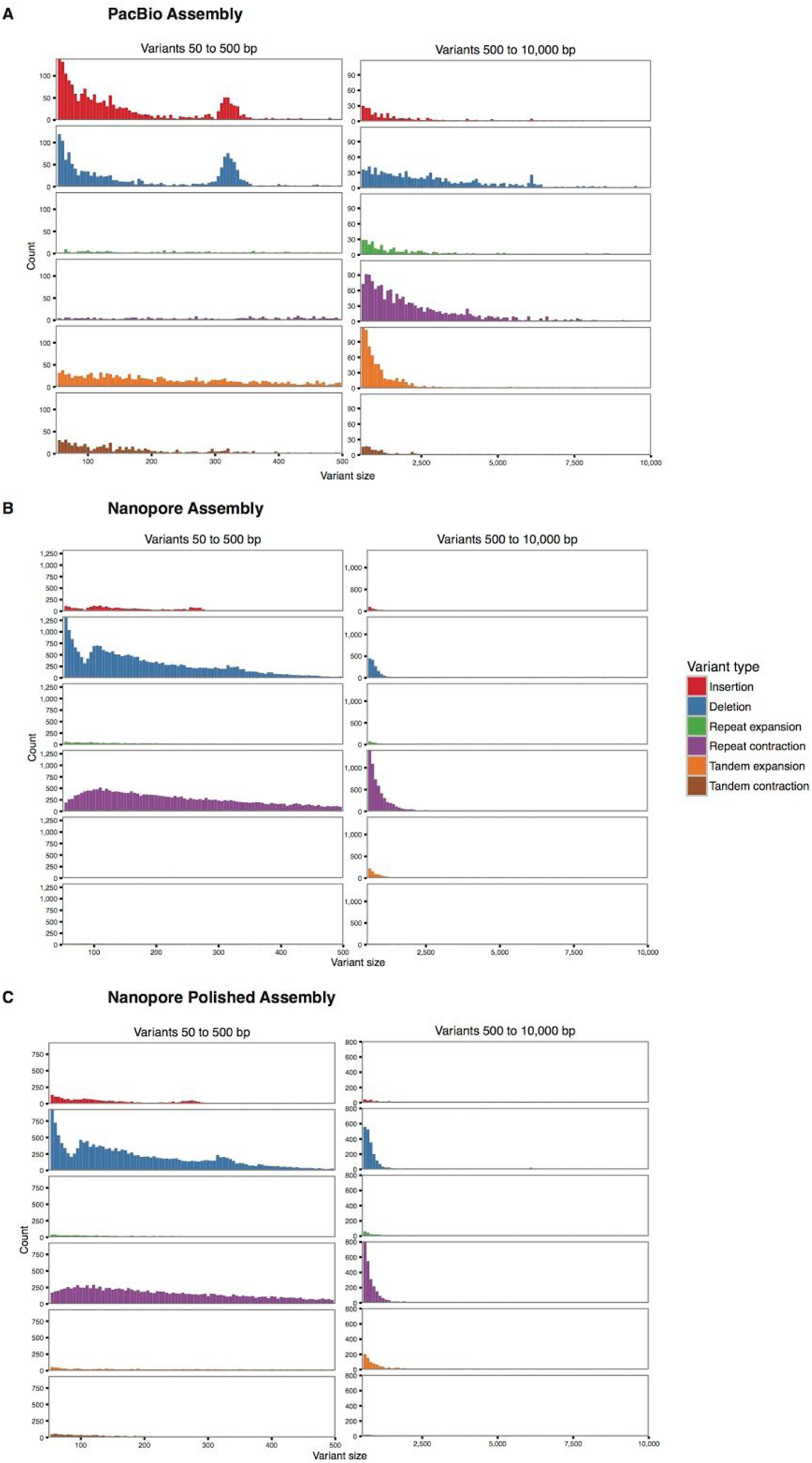
Structural Variant Analysis. Structural variants in the whole-genome nanopore assembly were identified using Assemblytics ^41^ and compared with a previous PacBio assembly ^5^. Histograms are given for insertion, deletion, repeat expansion/contraction, and tandem expansion/contraction SVs versus GRCh38. These are further broken into small (50–500 bp) and large (500–10000 bp) categories. Notably, the PacBio assembly shows a balanced rate of insertions and deletions, with a peak at 300 bp due to Alu insertion and deletion. In contrast, the nanopore assembly shows a strong deletion bias, with the majority of variants being deletions <500 bp. Note that this changes the y-axis scale and obscures the Alu peaks in these plots. Post-polishing, the deletion bias is reduced but is still significantly higher than PacBio. It is expected that assembly of Scrappie reads would further reduce the deletion bias observed.

**SI Figure 6.**
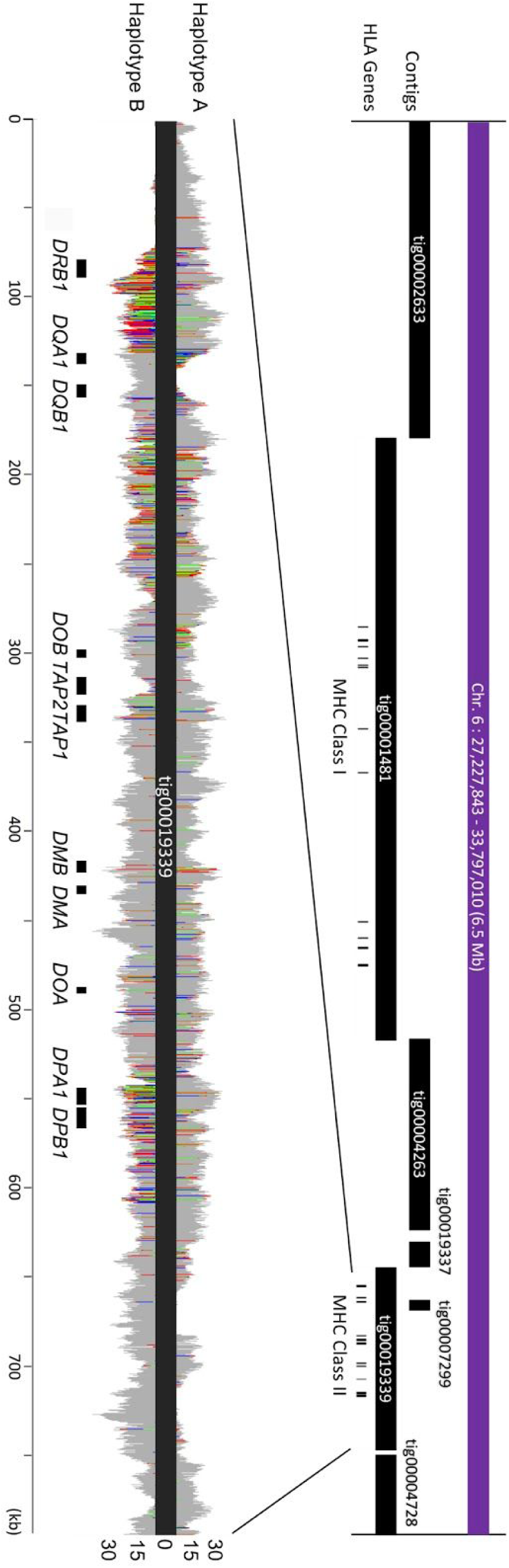
MHC Region. Canu contigs from the 30× nanopore assembly arranged along the MHC region of human chromosome 6 with HLA genes marked underneath as black bars. The zoomed region shows contig ‘tig00019339’ with corresponding nanopore read alignments separated by haplotype. Variants in the nanopore reads relative to the contig are shown as stacked and colored bars, as with IGV. Haplotype switching within the Canu contig is evident, but phasing was possible after assembly. Heterozygous variants in this contig were called from Illumina data and phased using nanopore reads with the WhatsHap program. Only reads with >1 phasing marker and >65% of variants in agreement are shown (79.3% of the total reads aligned to tig00019339).

**SI Figure 7.**
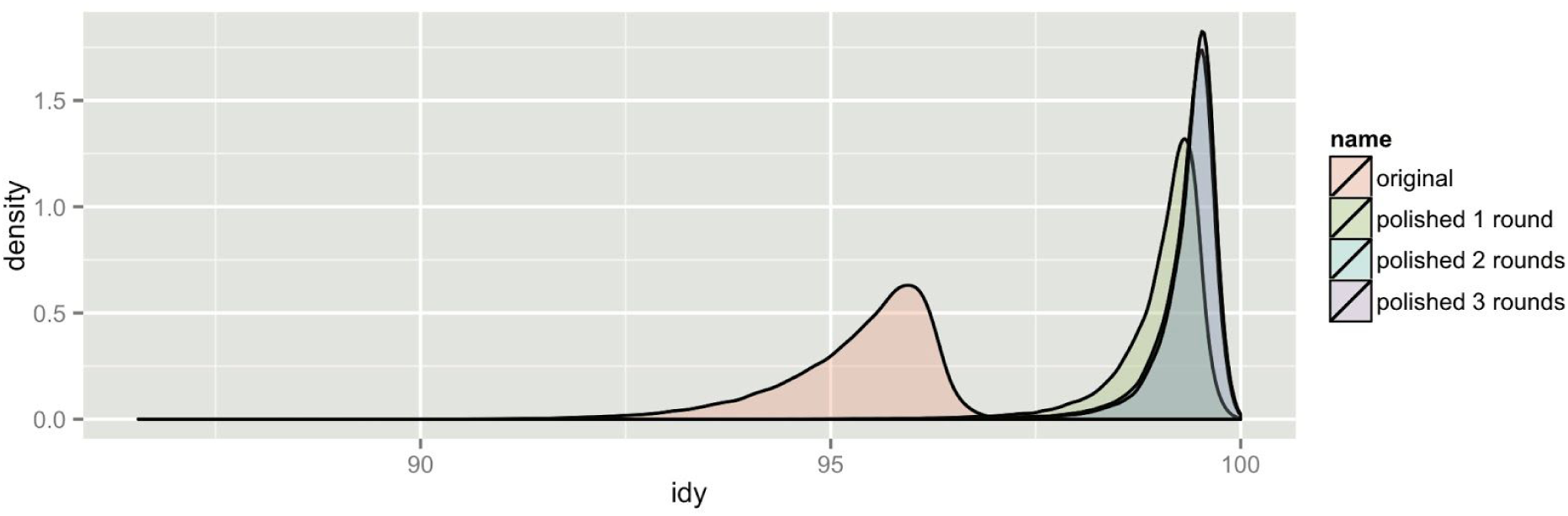
Assembly accuracy. Accuracy of the 30× nanopore assembly before and after Illumina polishing. Modal accuracy of the nanopore-only assembly is ∼96%. After Illumina polishing, this increases to >99%, with no substantial gain after 2 rounds of polishing.

**SI Figure 8.**
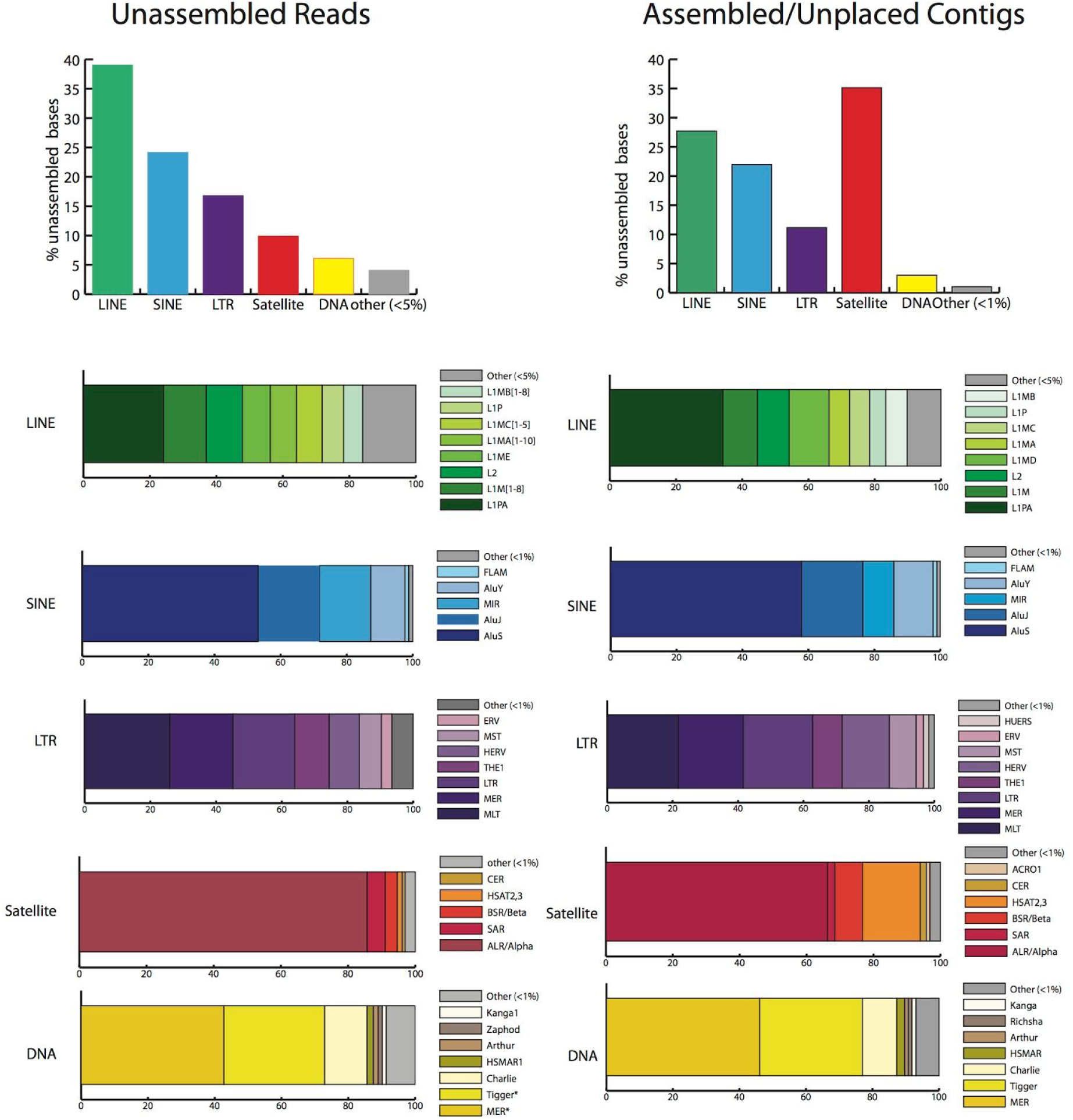
Sequences not found in the assembly. Distribution of repeat classes observed in unassembled sequence reads and contigs that were not incorporated in primary assembly. Percentage of bases for each repeat class are listed for both unassembled reads and assembled, yet unplaced contigs. Proportion of repeat families within the general repeat class are provided using sequence annotation by RepeatMasker (RepBase22.03).

**SI Figure 9.**
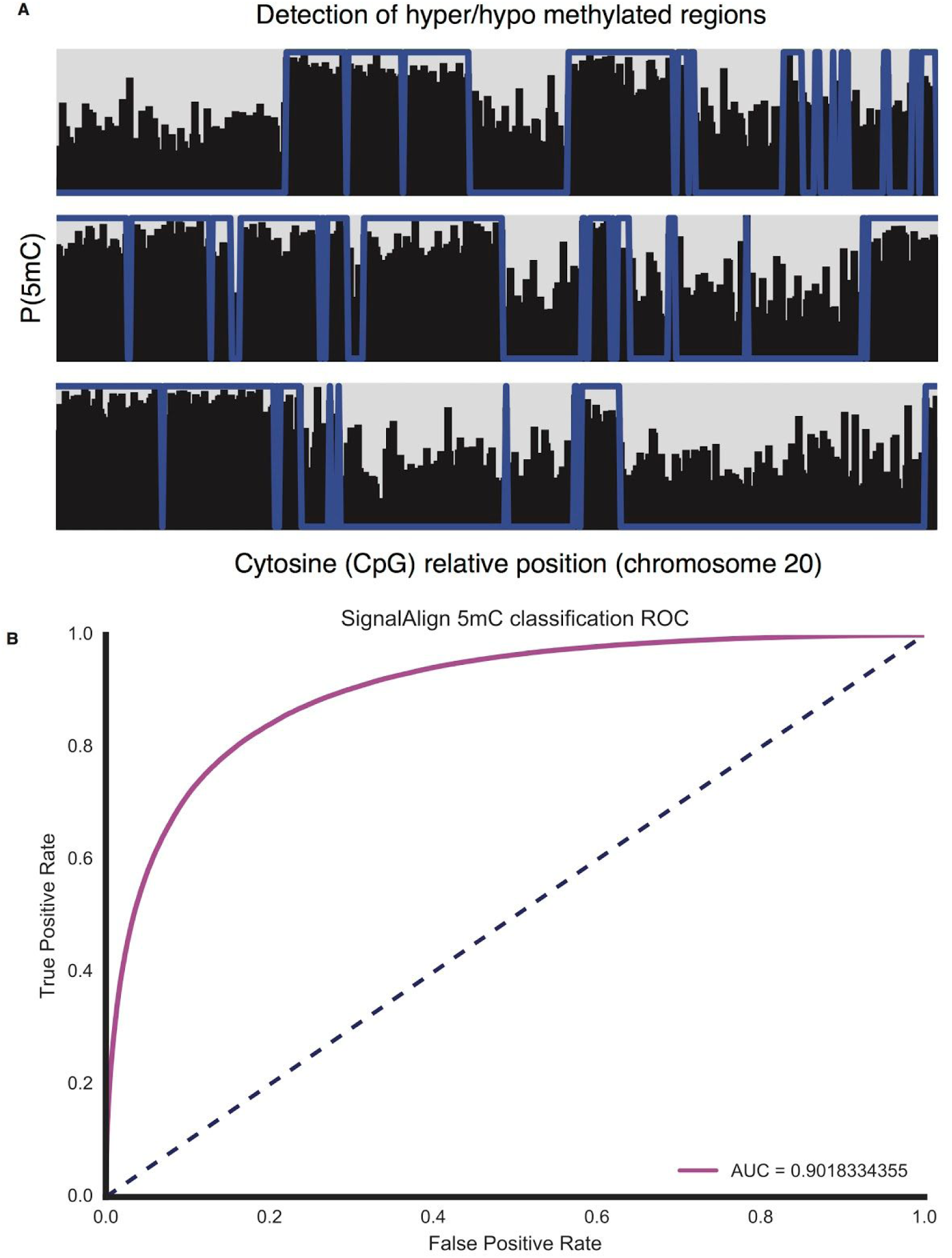
Methylation. A) Native DNA methylation detection on a selected portion of chromosome 20. Individual plots show 500 called cytosine bases ordered along chromosome 20. Total marginal probability of methylation is shown as black bar. High-confidence methylation calls from ENCODE (ENCSR890UQO), blue line, were filtered for positions where all reads called methylated or not methylation to remove ambiguity. Cytosine calls were filtered to only sites with coverage >= 10 reads in both data sets. B) Receiver operating characteristic (ROC) plot describing SignalAlign as a binary classifier for individual 5-methyl cytosine detection.

**SI Figure 10.**
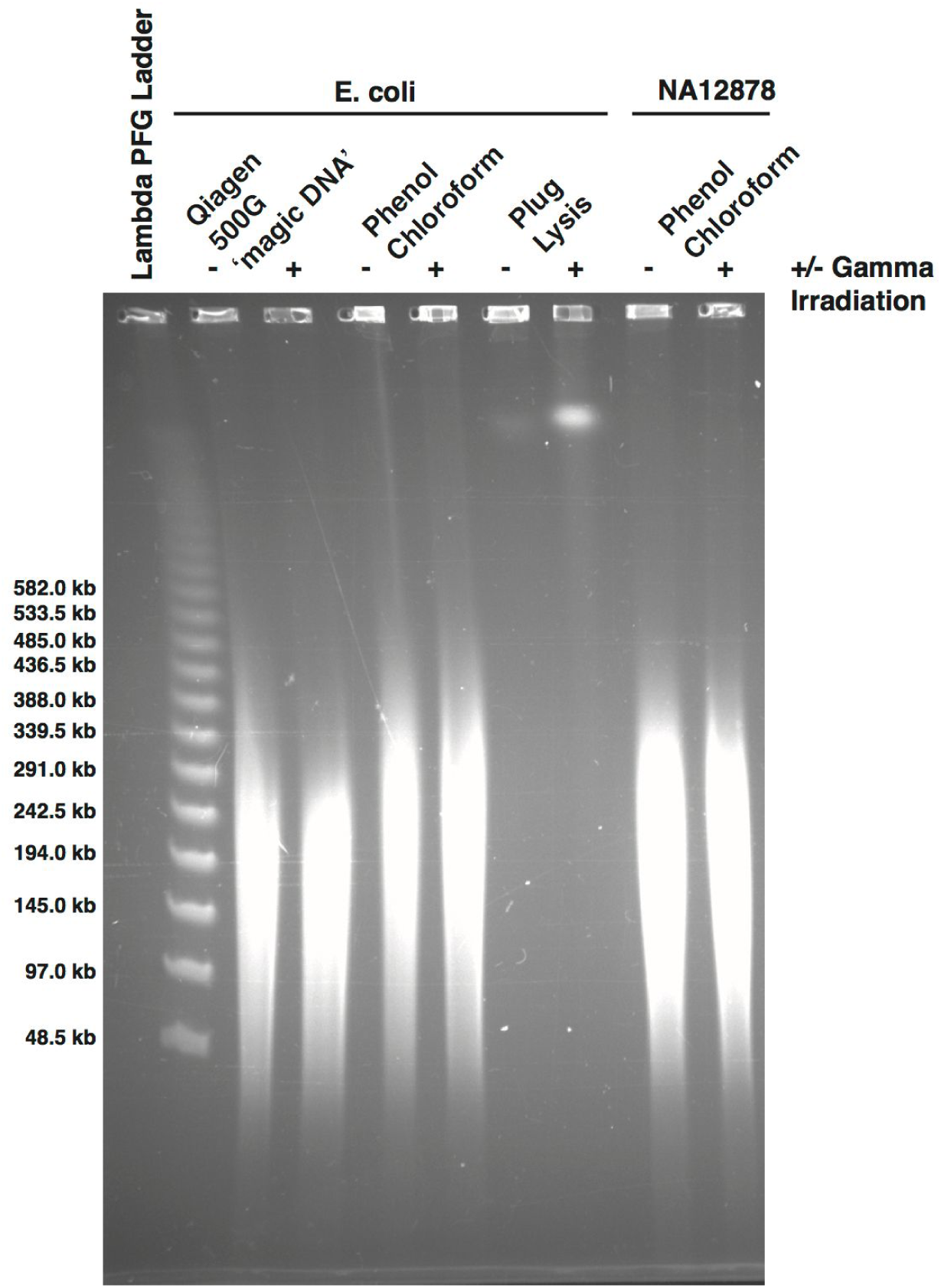
Ultra-long reads DNA extraction. Pulsed-field gel showing fragment sizes of; *E. coli* MG1655 DNA extracted using the Qiagen Genomic DNA buffer set and a Qiagen 500G column following the protocol for bacteria (lanes 2 and 3), E. coli MG1655 DNA extracted using the Sambrook and Russell phenol/chloroform protocol described in the methods section (lanes 4 and 5), E. coli MG1655 DNA extracted using a plug lysis method to preserve intact chromosomes (lanes 6 and 7) and Human NA12878 DNA extracted using the Sambrook and Russell phenol/chloroform protocol described in the methods section (lane 8 and 9). For each pair of samples one was irradiated with approximately 35 Gray ionising radiation to introduce double-strand breaks, this improves the intensity of the band representing the 4.6 Mb *E. coli* MG1655 chromosome. A 1.2% PFG agarose gel made with 0.5% TBE and run on a Bio-Rad CHEF Mapper at 14°C for 20 hours 46 minutes with a two-state 120° included angle, 6 V/cm gradient, initial switch time 0.64s and final switch time 13m 13.22s. The gel was ethidium bromide stained and imaged on a Bio-Rad Gel Doc XR system.

**SI Figure 11.**
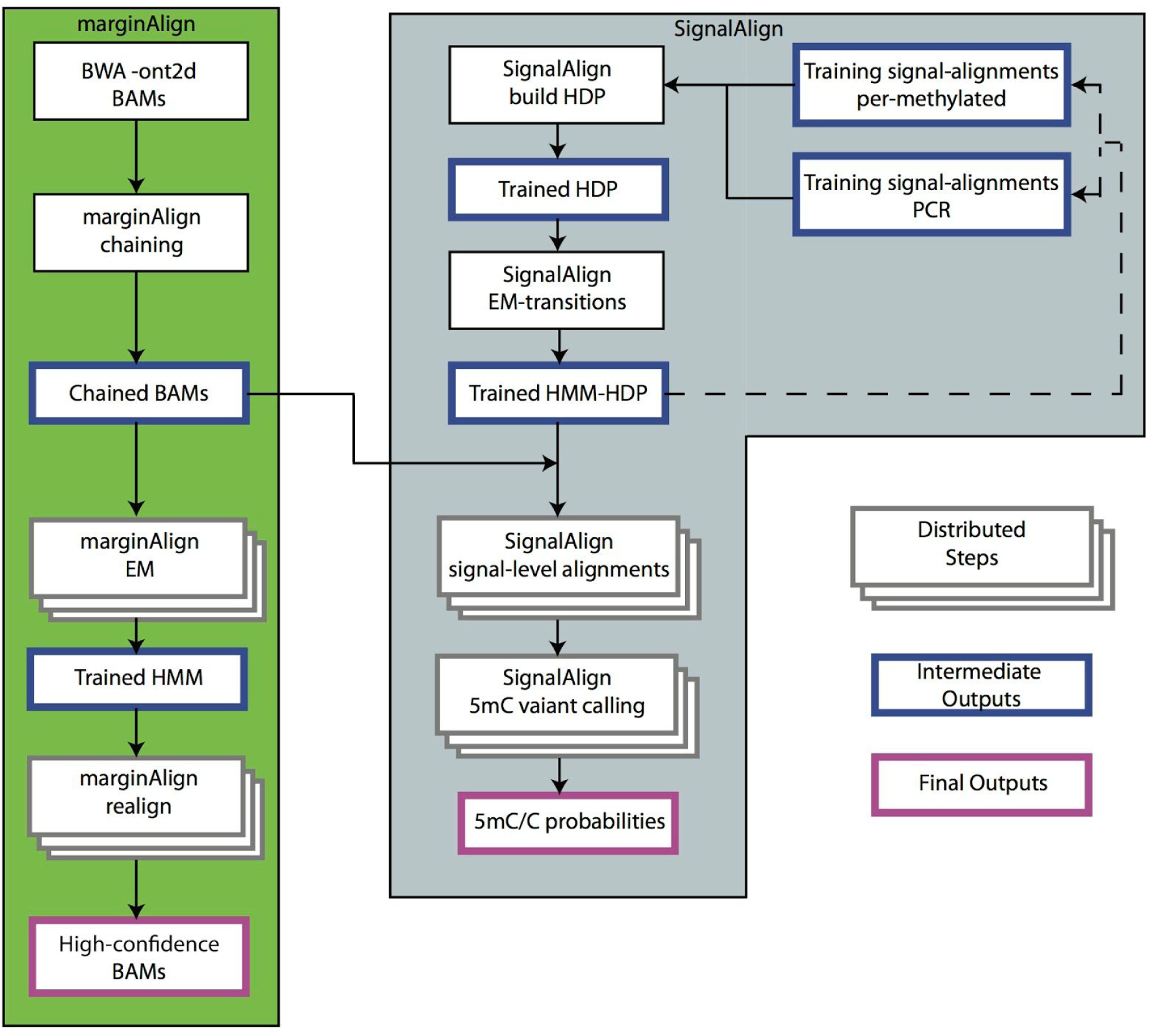
marginAlign/SignalAlign Work Flow. Workflow chart describing marginAlign and SignalAlign. All distributed steps were implemented as part of a Toil-pipeline to be run in the cloud. Dotted lines represent repeating steps (iterations).

### Supplementary Tables

**SI Table 1.** Summary sequencing statistics. Summary statistics for every flowcell used in this study.

| Flowcell ID | Number of Reads | Bases Sequenced | Date | Centre | Sample Type | Kit | Pore |
| --- | --- | --- | --- | --- | --- | --- | --- |
| FAB23716 | 356209 | 1409812422 | 14/07/16 | UBC | DNA | Rapid | R9 |
| FAB39088 | 658224 | 3287994454 | 19/09/16 | Notts | DNA | Ligation | R9.4 |
| FAB39075 | 466329 | 2439355478 | 20/09/16 | UBC | DNA | Ligation | R9.4 |
| FAB39043 | 436976 | 2273008592 | 23/09/16 | Bham | DNA | Ligation | R9.4 |
| FAB42706 | 430660 | 1966505502 | 12/10/16 | UBC | DNA | Ligation | R9.4 |
| FAB41174 | 117057 | 687394987 | 13/10/16 | Bham | DNA | Ligation | R9.4 |
| FAB42260 | 267644 | 1399557161 | 13/10/16 | UBC | DNA | Ligation | R9.4 |
| FAB42316 | 572838 | 3275026637 | 14/10/16 | Notts | DNA | Ligation | R9.4 |
| FAB42205 | 317654 | 1686630108 | 14/10/16 | Notts | DNA | Ligation | R9.4 |
| FAB42804 | 16669 | 75062609 | 14/10/16 | Bham | DNA | Ligation | R9.4 |
| FAB42561 | 233678 | 1520513556 | 19/10/16 | Notts | DNA | Ligation | R9.4 |
| FAB42473 | 644869 | 3357548938 | 19/10/16 | UBC | DNA | Ligation | R9.4 |
| FAB42395 | 38291 | 179704035 | 20/10/16 | Norwich | DNA | Ligation | R9.4 |
| FAB42476 | 435158 | 2363036522 | 27/10/16 | UBC | DNA | Ligation | R9.4 |
| FAB42451 | 817629 | 4530477841 | 28/10/16 | Notts | DNA | Ligation | R9.4 |
| FAB42704 | 276152 | 1750149482 | 28/10/16 | UBC | DNA | Ligation | R9.4 |
| FAB42828 | 33527 | 163405138 | 01/11/16 | Norwich | DNA | Ligation | R9.4 |
| FAB42810 | 322058 | 2020615256 | 02/11/16 | Norwich | DNA | Ligation | R9.4 |
| FAB42798 | 193551 | 1339441522 | 03/11/16 | Norwich | DNA | Ligation | R9.4 |
| FAB45280 | 128234 | 799554798 | 11/11/16 | Norwich | DNA | Ligation | R9.4 |
| FAB46664 | 491346 | 2038018797 | 15/11/16 | UBC | DNA | Ligation | R9.4 |
| FAB46683 | 72605 | 286275511 | 17/11/16 | Bham | DNA | Ligation | R9.4 |
| FAB45332 | 530938 | 2864140853 | 17/11/16 | UBC | DNA | Ligation | R9.4 |
| FAB44989 | 558224 | 3443824633 | 18/11/16 | UCSC | DNA | Ligation | R9.4 |
| FAB43577 | 426941 | 2539015084 | 18/11/16 | UCSC | DNA | Ligation | R9.4 |
| FAB45321 | 299174 | 2584017112 | 22/11/16 | Notts | Cells | Ligation | R9.4 |
| FAF01441 | 254705 | 2203636947 | 22/11/16 | Bham | Cells | Ligation | R9.4 |
| FAF01169 | 339447 | 2913892142 | 22/11/16 | Bham | Cells | Ligation | R9.4 |
| FAB45277 | 53547 | 445641679 | 22/11/16 | Notts | Cells | Ligation | R9.4 |
| FAF01127 | 632728 | 4972081712 | 25/11/16 | Bham | Cells | Ligation | R9.4 |
| FAF01132 | 689781 | 5455971336 | 25/11/16 | Bham | Cells | Ligation | R9.4 |
| FAB45271 | 472656 | 3689043164 | 28/11/16 | Notts | Cells | Ligation | R9.4 |
| FAB45321 | 123037 | 1043504055 | 28/11/16 | Notts | Cells | Ligation | R9.4 |
| FAB49914 | 309175 | 2841008085 | 28/11/16 | Notts | Cells | Ligation | R9.4 |
| FAB49712 | 632158 | 4906148911 | 28/11/16 | Bham | Cells | Ligation | R9.4 |
| FAF01253 | 471698 | 3695661984 | 28/11/16 | Bham | Cells | Ligation | R9.4 |
| FAB49164 | 746333 | 4438258089 | 06/12/16 | UCSC | DNA | Ligation | R9.4 |
| FAF04090 | 91304 | 1213584440 | 09/12/16 | Bham | Cells | Rapid | R9.4 |
| FAB49908 | 224380 | 3141600861 | 09/12/16 | Bham | Cells | Rapid | R9.4 |
| FAF15665 | 82138 | 1806857522 | 10/03/17 | Notts | Cells | Ultrareads | R9.4 |
| FAF13748 | 53723 | 1252868852 | 10/03/17 | Notts | Cells | Ultrareads | R9.4 |
| FAF10039 | 41385 | 848632752 | 01/03/17 | Bham | Cells | Ultrareads | R9.4 |
| FAF09968 | 19674 | 594496244 | 03/03/17 | Bham | Cells | Ultrareads | R9.4 |
| FAF09277 | 73755 | 1987434656 | 03/06/17 | Bham | Cells | Ultrareads | R9.4 |
| FAF14035 | 75692 | 1831031405 | 08/03/17 | Notts | Cells | Ultrareads | R9.4 |
| FAF15694 | 61227 | 1533616061 | 06/03/17 | Bham | Cells | Ultrareads | R9.4 |
| FAF09713 | 65142 | 1639658993 | 07/03/17 | Bham | Cells | Ultrareads | R9.4 |
| FAF18554 | 270189 | 2730589684 | 06/03/17 | UBC | Cells | Rapid | R9.4 |
| FAF15630 | 9663 | 322753214 | 09/03/17 | Notts | Cells | Ultrareads | R9.4 |
| FAF09640 | 72936 | 1496943560 | 07/03/17 | Bham | Cells | Ultrareads | R9.4 |
| FAF09701 | 68169 | 1731054841 | 03/03/17 | Bham | Cells | Ultrareads | R9.4 |
| FAF15586 | 71155 | 1750584936 | 08/03/17 | Bham | Cells | Ultraread | R9.4 |
| FAF05869 | 451020 | 3613667827 | 08/03/17 | UBC | Cells | Ligation | R9.4 |

**SI Table 2.** Summary alignment statistics by Flowcell. Summary alignment statistics for each flow cell, excluding ultra-long reads.

| Flow Cell ID | Number of Sequences | Mapped Reads | Mapped MQ0 | Unmapped | Bases Mapped | Avg Length |
| --- | --- | --- | --- | --- | --- | --- |
| FAB23716 | 356209 | 319259 | 26702 | 36950 | 1165998694 | 3957 |
| FAB39088 | 658224 | 613044 | 35394 | 45180 | 3007307322 | 4995 |
| FAB39075 | 466329 | 425117 | 28167 | 41212 | 2146453407 | 5230 |
| FAB39043 | 436976 | 415389 | 21043 | 21587 | 2113140439 | 5201 |
| FAB42706 | 430660 | 375374 | 17378 | 55286 | 1867123361 | 4566 |
| FAB41174 | 117057 | 114520 | 4186 | 2537 | 652217119 | 5872 |
| FAB42260 | 267644 | 246982 | 15624 | 20662 | 1263089767 | 5229 |
| FAB42804 | 16669 | 13311 | 1755 | 3358 | 53666089 | 4503 |
| FAB42316 | 572838 | 512994 | 18985 | 59844 | 3100596254 | 5717 |
| FAB42205 | 317654 | 282502 | 12561 | 35152 | 1601397762 | 5309 |
| FAB42561 | 233678 | 225141 | 10255 | 8537 | 1420740185 | 6506 |
| FAB42473 | 644869 | 611138 | 32539 | 33731 | 3112342902 | 5206 |
| FAB42395 | 38291 | 36477 | 2059 | 1814 | 167168840 | 4693 |
| FAB42476 | 435158 | 416969 | 20908 | 18189 | 2214880871 | 5430 |
| FAB42451 | 817629 | 779328 | 36986 | 38301 | 4178966543 | 5540 |
| FAB42704 | 276152 | 263722 | 12926 | 12430 | 1619875186 | 6337 |
| FAB42828 | 33527 | 27843 | 2442 | 5684 | 146819837 | 4873 |
| FAB42810 | 322058 | 305070 | 16802 | 16988 | 1808343119 | 6274 |
| FAB42798 | 193551 | 185739 | 8749 | 7812 | 1232035338 | 6920 |
| FAB45280 | 128234 | 122219 | 6336 | 6015 | 743280816 | 6235 |
| FAB46664 | 491346 | 456247 | 27622 | 35099 | 1862427349 | 4147 |
| FAB46683 | 72605 | 64739 | 5307 | 7866 | 269213160 | 3942 |
| FAB45332 | 530938 | 497862 | 26392 | 33076 | 2620752139 | 5394 |
| FAB43577 | 426941 | 410137 | 19835 | 16804 | 2344990054 | 5946 |
| FAB44989 | 558224 | 536572 | 25936 | 21652 | 3161900821 | 6169 |
| FAF01169 | 339447 | 315489 | 16481 | 23958 | 2677881316 | 8584 |
| FAF01441 | 254705 | 238834 | 12458 | 15871 | 2010117898 | 8651 |
| FAB45277 | 53547 | 51957 | 2132 | 1590 | 426639054 | 8322 |
| FAB45321 | 299174 | 283355 | 15165 | 15819 | 2366003310 | 8637 |
| FAF01127 | 632728 | 605633 | 27192 | 27095 | 4640355789 | 7858 |
| FAF01132 | 689781 | 655357 | 33564 | 34424 | 4966810089 | 7909 |
| FAB49712 | 632158 | 612752 | 26264 | 19406 | 4594356245 | 7760 |
| FAF01253 | 471698 | 454434 | 20639 | 17264 | 3430678969 | 7834 |
| FAB45321 | 123037 | 118311 | 5891 | 4726 | 952851126 | 8481 |
| FAB49914 | 309175 | 296250 | 12281 | 12925 | 2673848960 | 9188 |
| FAB45271 | 472656 | 450702 | 20148 | 21954 | 3468377327 | 7804 |
| FAB49164 | 746333 | 718351 | 32664 | 27982 | 4107087899 | 5946 |
| FAB49908 | 224380 | 211060 | 11903 | 13320 | 2898563539 | 14001 |
| FAF04090 | 91304 | 83164 | 6072 | 8140 | 1085757398 | 13291 |

**SI Table 3.** Summary alignment statistics by chromosome. (Excluding ultra-long reads)

| Chromosome | Number of Mapped Reads | Number of Mapped Reads (MQ0) | Number of Bases Mapped | Avg Length |
| --- | --- | --- | --- | --- |
| Chr 1 | 1075867 | 43397 | 6829526262 | 6744 |
| Chr 2 | 1062314 | 31802 | 6755642896 | 6842 |
| Chr 3 | 858643 | 24189 | 5487703898 | 6757 |
| Chr 4 | 845677 | 30723 | 5395140705 | 6890 |
| Chr 5 | 774613 | 23499 | 4953273570 | 6821 |
| Chr 6 | 723047 | 24496 | 4618883250 | 6762 |
| Chr 7 | 696473 | 28231 | 4382999832 | 6772 |
| Chr 8 | 617988 | 23361 | 3968911801 | 6844 |
| Chr 9 | 539660 | 25898 | 3428430670 | 6764 |
| Chr 10 | 594688 | 20787 | 3805443564 | 6845 |
| Chr 11 | 583055 | 17748 | 3710684724 | 6855 |
| Chr 12 | 586663 | 17891 | 3734922623 | 6840 |
| Chr 13 | 440615 | 17662 | 2844212242 | 6904 |
| Chr 14 | 383777 | 15752 | 2439119767 | 6713 |
| Chr 15 | 359853 | 19556 | 2268233023 | 6838 |
| Chr 16 | 386401 | 22680 | 2425913744 | 6787 |
| Chr 17 | 369036 | 22907 | 2302471086 | 6661 |
| Chr 18 | 339094 | 13053 | 2172098564 | 6807 |
| Chr 19 | 257039 | 10926 | 1472760724 | 6266 |
| Chr 20 | 291960 | 13226 | 1829244829 | 6659 |
| Chr 21 | 192383 | 24988 | 1207807437 | 6792 |
| Chr 22 | 172934 | 10514 | 1041347396 | 6665 |
| Chr X | 658347 | 28769 | 4210769167 | 7076 |
| Chr Y | 23378 | 5292 | 133803203 | 7869 |
| Chr M | 59363 | 658 | 91949786 | 1628 |

**SI Table 4.** Read Length Metrics by DNA Input. Summary read length metrics subdivided by DNA preparation and sequencing library preparation method.

| **Input DNA** | **Number of Reads** | **Median Read Length** | **Mean Read Length** | **Total Bases** | **Read N50** |
| --- | --- | --- | --- | --- | --- |
| Ligation Library, Cell DNA | 4278106 | 7140 | 8123 | 34750607127 | 12112 |
| Ligation Library, Cell DNA (Albacore/MinKNOW 1.4 Control) | 451020 | 4463 | 8012 | 3613667827 | 13920 |
| Ligation Library, Coriell DNA | 9233585 | 3853 | 5493 | 50724515583 | 9136 |
| Rapid Kit, Cell DNA | 315684 | 6800 | 13796 | 4355185301 | 30397 |
| Rapid Kit, Cell DNA (Albacore/MinKNOW 1.4 Control) | 270189 | 4073 | 10106 | 2730589684 | 24848 |
| Rapid Kit, Coriell DNA | 356209 | 2375 | 3958 | 1409812422 | 6978 |
| Ultralong reads Protocol, Cell DNA | 694659 | 3488 | 24179 | 16795933036 | 99790 |

**SI Table 5.** Summary containing kmer counts with respect to chromosome 20 for each of the base callers used in this study: See Supplementary_Table5_kmer.xlsx

**SI Table 6.** Structural Variants. Summary of structural variants observed in the Nanopore only and Nanopore polished canu assemblies.

|  |  | PacBio |  | Nanopore |  | Nanopore Polished |  |
| --- | --- | --- | --- | --- | --- | --- | --- |
| SV Type | Size Range (bp) | Count | Total bp | Count | Total bp | Count | Total bp |
| Insertion | 50-500 | 2042 | 306272 | 3365 | 610141 | 2446 | 383069 |
|  | 500-10000 | 319 | 618769 | 535 | 767711 | 394 | 681803 |
|  | **Total:** | 2361 | 925041 | 3900 | 1377852 | 2840 | 1064872 |
| Deletion | 50-500 | 1645 | 290106 | 26853 | 4718511 | 18359 | 3358506 |
|  | 500-10000 | 1079 | 3142086 | 1938 | 2047493 | 2438 | 2528925 |
|  | **Total:** | 2724 | 3432192 | 28791 | 6766004 | 20797 | 5887431 |
| Tandem expansion | 50-500 | 1431 | 316231 | 1199 | 358827 | 1782 | 422450 |
|  | 500-10000 | 778 | 1024584 | 1067 | 1216138 | 1068 | 1285297 |
|  | **Total:** | 2209 | 1340815 | 2266 | 1574965 | 2850 | 1707747 |
| Tandem contraction | 50-500 | 685 | 114672 | 629 | 96955 | 1286 | 209359 |
|  | 500-10000 | 159 | 251549 | 75 | 158553 | 186 | 345439 |
|  | **Total:** | 844 | 366221 | 704 | 255508 | 1472 | 554798 |
| Repeat expansion | 50-500 | 241 | 57575 | 1961 | 389003 | 1289 | 251816 |
|  | 500-10000 | 373 | 894444 | 469 | 688134 | 420 | 707130 |
|  | **Total:** | 614 | 952019 | 2430 | 1077137 | 1709 | 958946 |
| Repeat contraction | 50-500 | 368 | 104240 | 24178 | 5467319 | 14276 | 3258222 |
|  | 500-10000 | 1627 | 3718349 | 6882 | 6930904 | 3129 | 2810950 |
|  | **Total:** | 1995 | 3822589 | 31060 | 12398223 | 17405 | 6069172 |
| **Totals for all variants:** |  | 10747 | 10838877 | 69151 | 23449689 | 47073 | 16242966 |

**SI Table 7.**
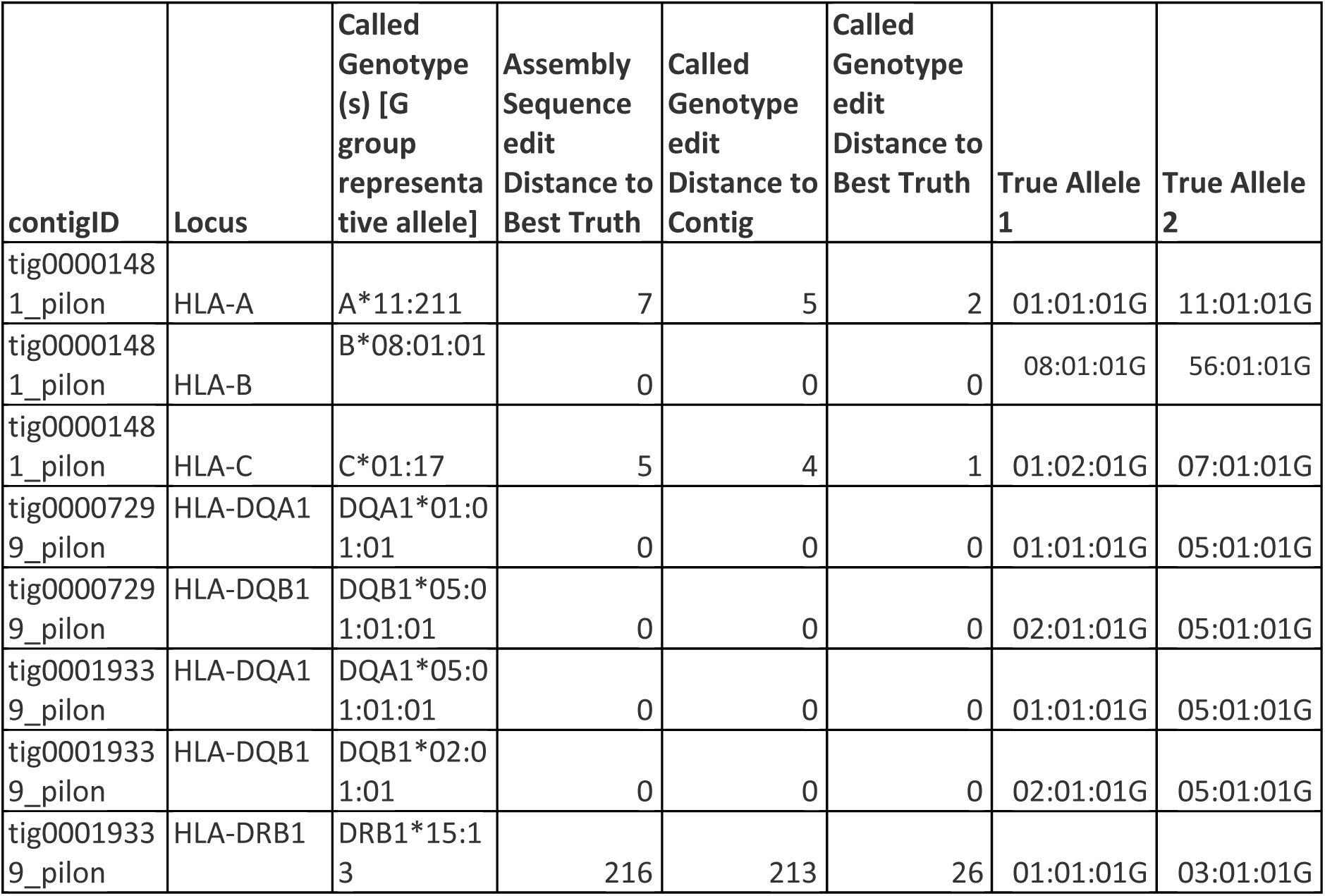
Summary of HLA Typing.

